# Identification of a Putative Metal Transporter in the Apicoplast of Malaria Parasites

**DOI:** 10.64898/2026.05.18.725985

**Authors:** Shai-anne Nalder, Adarsh K. Mayank, Willisa Liou, James A. Wohlschlegel, Paul A. Sigala

## Abstract

*Plasmodium falciparum* malaria parasites harbor an essential plastid organelle, called the apicoplast, which produces key metabolites required for organelle function and parasite viability. Apicoplast functions depend on iron and other metals, but the membrane transporters that mediate metal import into this organelle have been challenging to identify. Tetracycline antibiotics, including doxycycline, specifically target the apicoplast and can exhibit metal-dependent activity. Using tetracycline-affinity proteomics, we identified a doxycycline-interacting, uncharacterized transmembrane protein (UCT) targeted to the apicoplast periphery but not proteolytically processed. Although lacking sequence similarity to proteins of known function, UCT has a predicted structure with high similarity to pentameric CorA-family metal transporters that mediate metal uptake in other organisms. Functional tests revealed that UCT is dispensable for blood-stage asexual parasites, suggesting that the apicoplast has evolved redundant mechanisms for metal uptake. UCT knockdown in gametocytes, however, impairs the development of sexual parasites, which are critical for mosquito transmission. Our study identifies an apicoplast membrane protein with localization and structural properties that predict a role in metal transport into this key organelle. This discovery can provide a biochemical springboard to unravel broader apicoplast mechanisms of metal uptake across multiple stages of parasite development, including mosquito-stage parasites that display heightened UCT expression.

## Introduction

Malaria is a deadly and ancient infectious disease that has plagued humankind for millennia. There was a steady decline in global malaria deaths since 2000, but this progress has now stalled, with ∼600,000 deaths annually since 2015^1^. *Plasmodium* parasites cause malaria, with most deaths resulting from *P. falciparum* infection. These single-celled eukaryotic organisms diverged from well-studied yeast and mammals early in eukaryotic evolution and acquired specialized adaptations to invade and replicate within iron-rich human erythrocytes. Identification and understanding of these molecular adaptations will give insight into the evolution of *Plasmodium* and other parasites in the phylum Apicomplexa, introduce new biological paradigms for key cellular processes, and reveal critical parasite vulnerabilities suitable for therapeutic targeting.

*Plasmodium* parasites harbor an essential and non-photosynthetic plastid organelle, called the apicoplast, that is derived from secondary endosymbiosis^2^ and thus surrounded by four membranes^3^ (Figure 1). Multiple metabolic pathways are targeted to this organelle, including the enzymes required for essential synthesis of the isoprenoid precursors, isopentenyl pyrophosphate (IPP) and dimethylallyl pyrophosphate. Several proteins involved in isoprenoid synthesis depend on metal cofactors, especially Fe-S clusters produced in the apicoplast by the SUF pathway^2^. IPP production depends on the ferredoxin/ferredoxin-NADP^+^ reductase (FD/FNR) system, which donates electrons to hydroxymethylbutenyl diphosphate synthase (IspG) and hydroxymethylbutenyl diphosphate reductase (IspH) that catalyze the final two steps of IPP production^4^. Both the FD/FNR system and the enzymes IspG and IspH require iron-sulfur clusters, making iron indispensable for IPP production and parasite viability^4^ (Figure 1).

**Figure 1.**
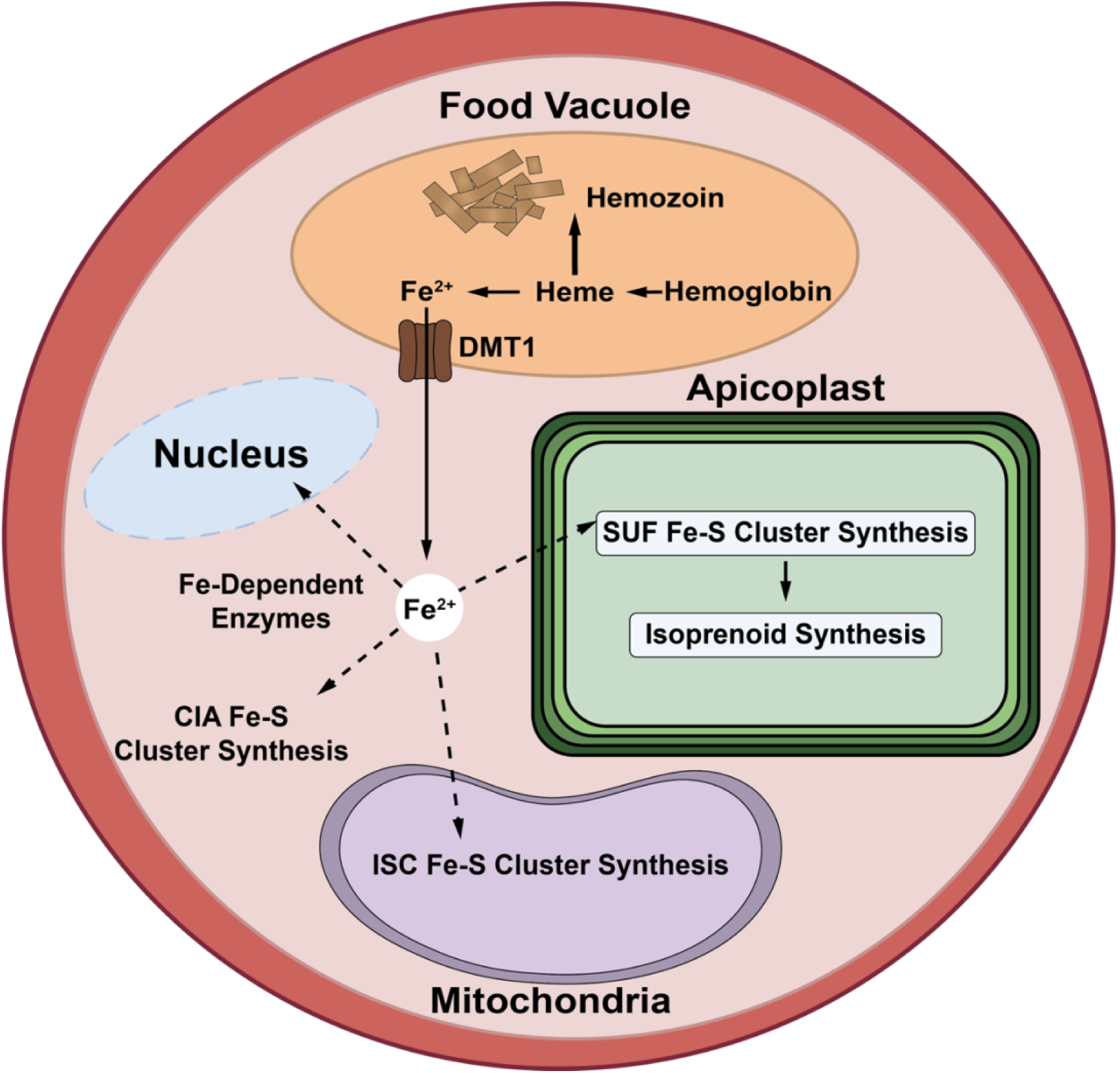
Scheme of iron mobilization in intraerythrocytic *P. falciparum* parasite depicting hemoglobin digestion to liberate free heme that is either biomineralized into inert hemozoin or proposed to be nonenzymatically degraded to produce labile iron. Labile iron is transported out of the food vacuole by DMT1, trafficked throughout the parasite via undefined mechanisms (dashed arrows), and delivered to support the iron-dependent pathways and enzymes in the apicoplast as well as mitochondrion, cytoplasm, and nucleus.

The primary source of iron within the parasite is likely the acidic food vacuole (FV), where the digestion of hemoglobin liberates free heme. Most of this heme is biomineralized into chemically inert hemozoin crystals^5^, but a trace amount is thought to undergo nonenzymatic degradation within the oxidizing FV environment to generate labile iron. Recent studies have provided evidence that this iron is transported out of the FV by PfDMT1 to supply nutritional iron to support broad parasite metabolism^6, 7^, including IPP synthesis and apicoplast biogenesis. Despite the central role of iron-dependent metabolism in apicoplast function, the mechanisms of iron uptake and internalization by the organelle remain elusive (Figure 1).

Metal uptake into the apicoplast is expected to require multiple distinct protein transporters with differing specificities to translocate iron and other metals across the four apicoplast membranes for utilization in the organelle matrix. Although substantial progress has been made in recent years to define the metabolic functions of the apicoplast and its metal requirements^8^, the identification of the apicoplast transporters responsible for importing iron and other key metals remains a significant knowledge gap in parasite biology^9^. Indeed, few metal transporters, and no specific iron transporters, have been identified in the apicoplast. Zip1 (Pf3D7_0609100) was identified as a putative zinc transporter with lower affinity for iron transport and was suggested to target the parasite plasma membrane and apicoplast^10^. However, multiple observations suggest that Zip1 does not play a central role in metal uptake into the apicoplast, including the proposed direction of Zip1 transport into the cytoplasm^10^ and the exclusive observation of Zip1 on the plasma membrane in a second study that also confirmed its non-essential function in blood-stage parasites^10, 11^.

Due to its prokaryotic ancestry, apicoplast metabolism is sensitive to several antibacterial translation inhibitors, including doxycycline (dox), which blocks apicoplast biogenesis at concentrations of 1-2 µM and kills parasites slowly over 96 hours^12–14^. In recent work, we discovered that higher-dose 10 µM dox kills parasites with faster, 48-hour activity through a distinct, apicoplast-specific mechanism that blocks organelle biogenesis^15^. This activity is selectively sensitive to exogenous iron but not to other metals, such as calcium, suggesting that 10 µM dox may interfere with apicoplast iron metabolism by an unknown mechanism. This observation, combined with the known propensity of dox and other tetracycline antibiotics to tightly bind metals^16^, suggested that these compounds might serve as chemical probes to identify proteins involved in metal metabolism and/or uptake into the apicoplast.

We used a tetracycline affinity resin to identify dox-specific interacting proteins in parasite lysates. Based on this approach, we identified an apicoplast-targeted protein (Pf3D7_1248300) that lacks sequence homology to proteins of known function but contains multiple transmembrane domains. Structural modeling predicted high structural similarity to metal transporters in bacteria and supported the formation of a multimeric pore similar to pentameric CorA-family transporters. We localized this protein to the apicoplast periphery and determined that it is not proteolytically processed, unlike most known apicoplast proteins. Functional studies suggest that UCT has a key role in the development of sexual gametocytes. This work unveils an apicoplast membrane protein with cryptic properties that suggest a critical function in metal uptake into this essential organelle and that can serve as a mechanistic foothold to unravel broader apicoplast import functions.

## Results

### Affinity Isolation Identifies a Doxycycline-Interacting Protein with Predicted Apicoplast Targeting

In bacteria, doxycycline binds to macromolecular targets that include the tetracycline repressor protein^17, 18^ and 30S ribosomes^19, 20^ via metal-dependent interactions. Our prior study revealed that 10 µM dox specifically inhibits apicoplast biogenesis over 48 hours via an iron-sensitive mechanism^15^. We therefore reasoned that metal-dependent activity of dox targeting the apicoplast might reflect interactions with one or more protein targets within this endosymbiotic organelle of cyanobacterial origin. To test this hypothesis, we carried out an affinity isolation experiment using solubilized parasite lysates and agarose beads covalently bound to tetracycline, a structural isomer of doxycycline that shares the same metal-binding sites^21^. To leverage tetracycline agarose to identify dox-interacting proteins, we performed an affinity pull-down of parasite lysates in the absence or presence of competing free dox. We also included a sample in which the parasite apicoplast was stably disrupted to assess similarities and differences in the interactome and the dependence of any affinity interactions on apicoplast status (Figure 2A). After washing and eluting the agarose beads with free dox, we used tandem mass spectrometry to identify proteins selectively enriched in the pull-down from untreated or apicoplast-disrupted lysates compared to dox-pretreated lysates.

**Figure 2.**
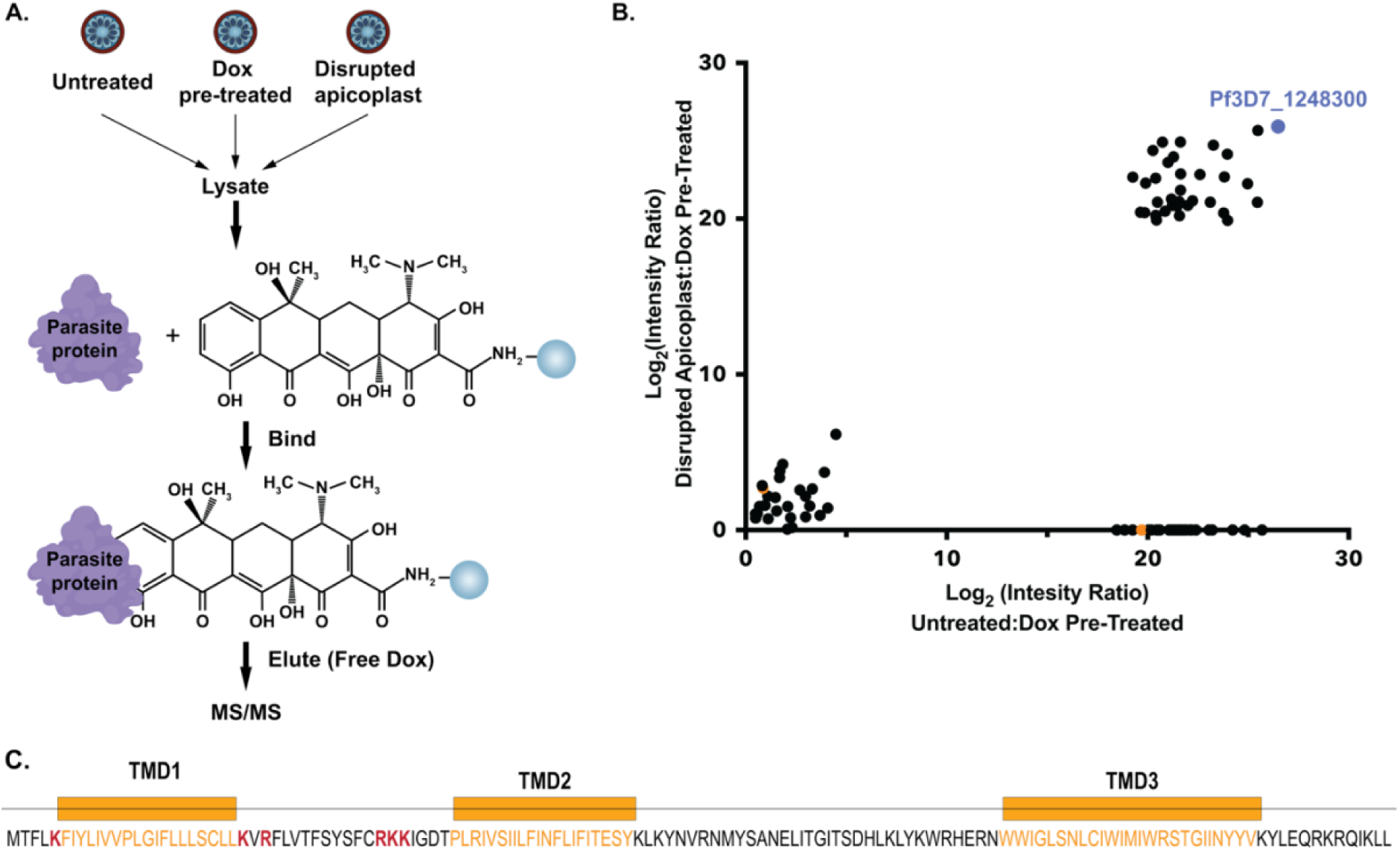
Affinity isolation identifies an uncharacterized apicoplast protein. (A) Affinity-isolation scheme showing untreated, dox pretreated, and disrupted-apicoplast lysates as inputs for affinity pulldown with tetracycline agarose. (B) Proteomic results showing the log_2_ ratio of spectral abundance for proteins identified in untreated versus dox-pretreated or disrupted apicoplast versus dox-pretreated lysates. Pf3D7_1248300 is shown in blue and non- or variably-enriched apicoplast proteins in orange (C) Protein map highlighting the three transmembrane domains (TMDs) in orange and positively charged residues (red) in the N-terminus of Pf3D7_1248300.

We identified a range of protein interactors but focused on the protein (Pf3D7_1248300) that was most highly enriched in both untreated and apicoplast-disrupted data sets compared to dox-pretreatment (Figure 2B, Figure S1, and Supplementary Data S1). This protein was previously identified in a curated apicoplast proteome^22^ based on proximity biotinylation, suggesting likely apicoplast targeting. We therefore focused on this protein and noted that its interaction with tetracycline agarose appeared to be independent of apicoplast status.

This 137-amino acid protein is conserved across *Plasmodium* species but lacks a predicted function based on its sequence dissimilarity to proteins of known function in other organisms. SignalP failed to identify an N-terminal signal peptide expected for targeting to the apicoplast. However, this protein has three predicted transmembrane domains, including one at the N-terminus that we hypothesized might function as a *de facto* signal peptide^23^. PlasmoAP also identified some features of an apicoplast transit peptide near the N-terminus that included multiple basic and positively charged residues^23^ (Figure 2C). These sequence features, combined with its prior reported detection in the apicoplast proteome, suggested possible but uncertain targeting of this protein to the apicoplast. We next carried out experiments to directly localize this protein in blood-stage parasites. For simplicity, we refer to this protein as an uncharacterized transmembrane protein, or UCT.

### UCT Localizes to the Apicoplast

Apicoplast-targeted proteins canonically require both an N-terminal signal peptide for entry into the endoplasmic reticulum (ER), followed by a positively charged transit peptide for routing to the apicoplast^23, 24^. Analysis of UCT utilizing the PlasmoAP too^l23^ suggested some features of a transit peptide, including positively charged clusters of N-terminal basic residues. However, it did not identify a canonical signal peptide. Instead, UCT has an N-terminal transmembrane domain (TMD) that may be sufficient to act as a signal peptide for ER entry^25^ (Figure 2C). Given prior detection of UCT in the apicoplast proteome^22^, we hypothesized that UCT is likely targeted to this organelle.

To directly localize UCT, we first stably transfected Dd2 parasites with a plasmid encoding expression of UCT with a C-terminal GFP fusion tag. The GFP signal observed in stably transfected live parasites appeared to branch and elongate throughout the parasite life cycle, which is typical of apicoplast proteins (Figure 3A and Figure S2A). To test apicoplast localization, we performed immunofluorescence analysis (IFA) and observed co-localization of UCT-GFP and the apicoplast acyl carrier protein (ACP) (Figure S2B).

**Figure 3.**
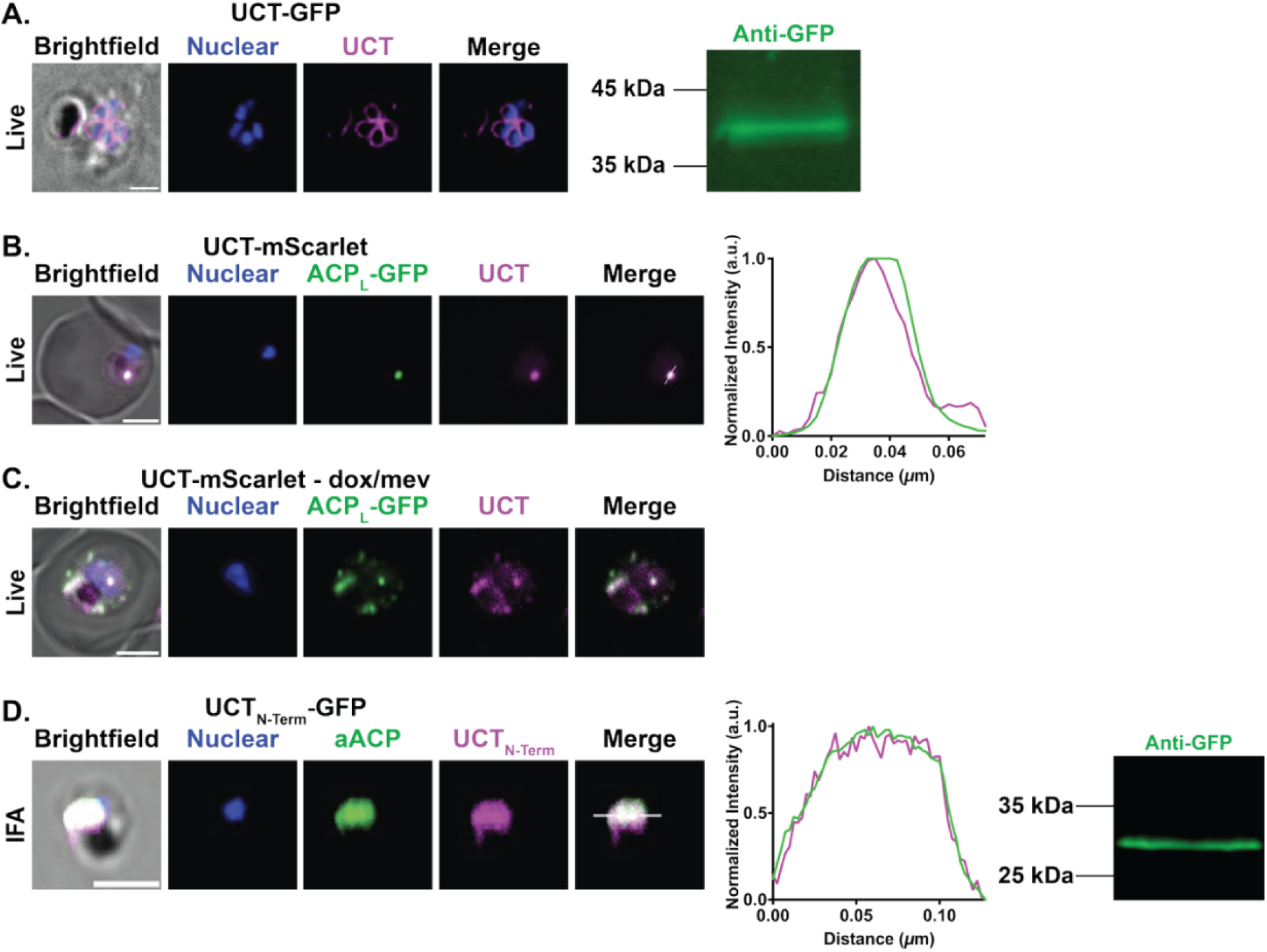
UCT localizes to the apicoplast. (A) Live-parasite fluorescent microscopy of UCT-GFP showing expected organelle branching, and anti-GFP western blot showing detection of UCT-GFP in lysates. (B) Live-parasite fluorescent miscopy of UCT-mScarlet (mS) showing co-localization with ACP_L_-GFP. (C) Live-parasite fluorescent microscopy of UCT-mS PfMev parasites treated with doxycycline and mevalonate showing dispersed foci of UCT-mS and ACP_L_-GFP upon apicoplast disruption. (D) Immunofluorescent microscopy images of (AA 1-42) UCT_N-term_-GFP showing co-localization with apicoplast ACP (aACP) and anti-GFP western blot of lysates. Scale bar = 2.5 microns.

To further confirm apicoplast localization of UCT, we transfected an episome encoding full-length UCT with a C-terminal mScarlet (mS) tag into NF54 PfMev parasites^26^, which express the ACP leader sequence (ACP_L_) fused to GFP as an apicoplast marker. Imaging live parasites by fluorescence microscopy revealed strong co-localization between UCT-mS and PfMev ACP_L_-GFP, with no substantial UCT signal beyond these co-localizing signals (Figure 3B and Figure S2C).

To further test that UCT targets the apicoplast, we stably disrupted the apicoplast by culturing parasites for multiple cycles in dox and mevalonate, which induces IPP synthesis in PfMev parasites and decouples parasite viability from apicoplast function^26^. Under these conditions, the apicoplast organelle is lost, and proteins that are targeted to the organelle are dispersed in cytoplasmic foci^14^. As expected for apicoplast targeting, UCT-mS and ACP_L_-GFP displayed a dispersed pattern of fluorescent foci in dox/mev-treated PfMev parasites (Figure 3C and Figure S2D). In addition, we stably disrupted the apicoplast in Dd2 UCT-GFP parasites and saw dispersed foci in live-cell imaging of GFP fluorescence, further confirming apicoplast targeting (Figure S2E).

To test whether the N-terminal region of UCT that includes the first TMD is sufficient to target GFP to the apicoplast, we stably transfected Dd2 parasites with an episome encoding the first 42 amino acids of UCT with a C-terminal GFP tag. IFA of fixed parasites revealed strong co-localization between UCT_N-term_-GFP and aACP, providing direct evidence that the N-terminus of UCT is sufficient for targeting to the apicoplast (Figure 3D and Figure S2E and H) and that the first TMD can function as a de facto signal peptide.

### UCT Lacks N-terminal Processing

Apicoplast-targeted proteins that are imported into the organelle matrix canonically display two bands by western blot that reflect proteolytic cleavage of the N-terminal transit peptide by the matrix-localized stromal processing peptidase, as previously shown for ACP^27^. However, there are several examples of apicoplast-targeted proteins that are not processed, including the outer triosephosphate transporter (PfoTPT, Pf3D7_0508300) that is targeted to the outer apicoplast membrane and displays only a single band via western blot that corresponds to the unprocessed protein^28^. Despite clear evidence for apicoplast localization of UCT-GFP, western blot analysis revealed a single ∼40 kDa band at the expected size for full-length UCT-GFP (Figure 3A and Figure S4A), suggesting a lack of processing. Western blot analysis of UCT_N-term_-GFP also revealed a single ∼30 kDa band at the expected size (Figure 3D and Figure S4B), further suggesting a lack of processing.

To compare the size of parasite-expressed UCT-GFP with the sizes expected for unprocessed or N-terminally processed proteins, we used western blot analysis to compare parental Dd2 parasites lacking tagged UCT, parasites expressing UCT-GFP that were untreated or treated for apicoplast disruption, and a co-load of untreated and disrupted parasites. We observed that disrupted and co-loaded samples co-migrated with untreated UCT-GFP, suggesting that UCT is not processed (Figure 4A and Figure S4A). To further test this result, we recombinantly expressed full-length UCT with a C-terminal GFP tag in *E. coli*, where no processing is expected. By western blot analysis, we observed that parasite-expressed UCT-GFP co-migrated with full-length bacterial UCT-GFP on SDS-PAGE (Figure 4B and Figure S4B), strongly suggesting that UCT is not processed upon apicoplast targeting. This result suggests that the UCT N-terminus is not exposed to the apicoplast matrix, possibly due to targeting to one of the three outer membranes of the apicoplast (Figure 1).

**Figure 4.**
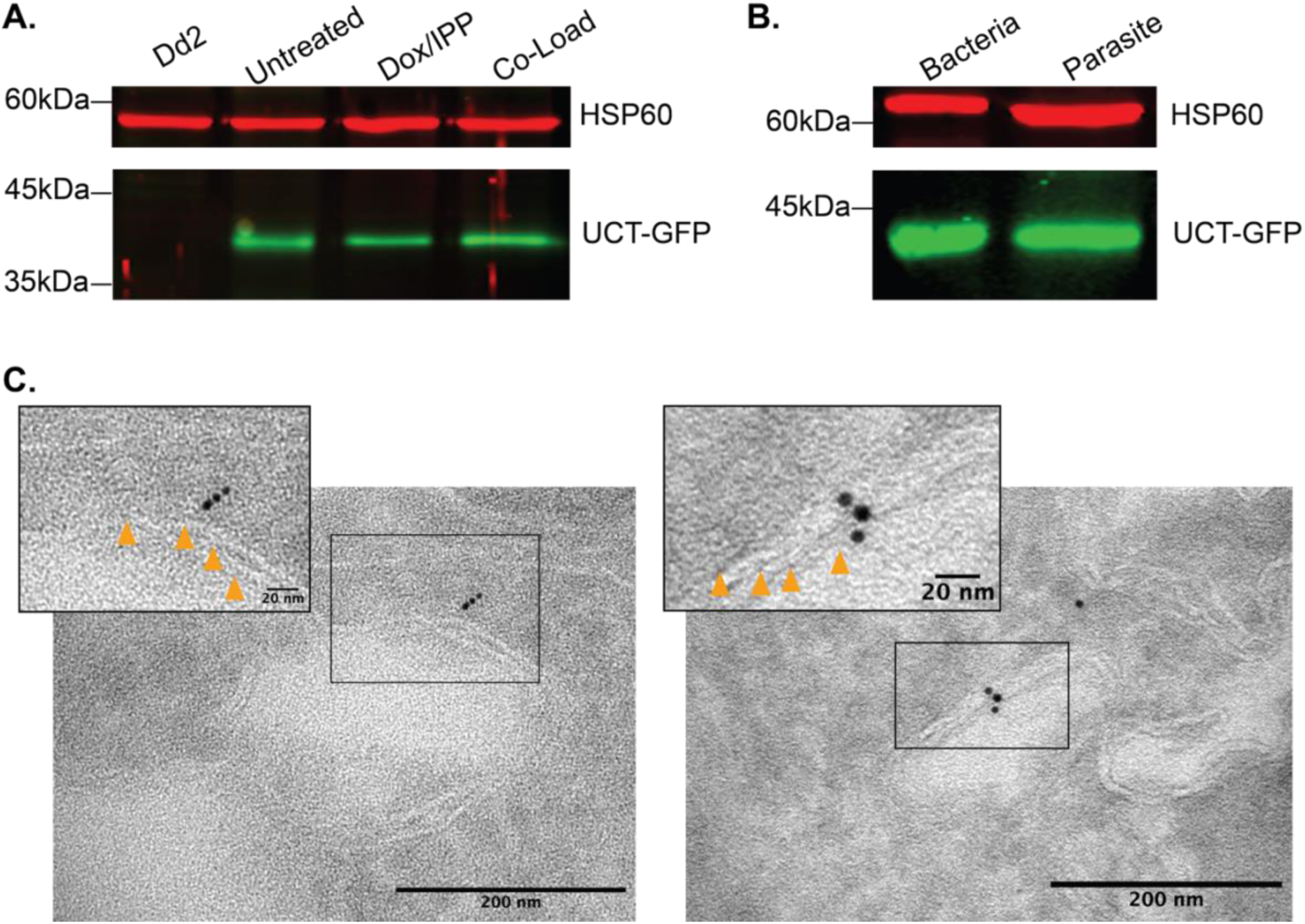
UCT is unprocessed and associated with apicoplast membranes. (A) Western blot of parental Dd2, untreated UCT-GFP, Dox/IPP disrupted UCT-GFP, and co-loaded untreated and disrupted parasites stained with anti-GFP and anti-HSP60 antibodies. (B) Western blot of full-length UCT-GFP recombinantly expressed in *E. coli* and UCT-GFP in parasites stained with anti-GFP and anti-HSP60 antibodies. (C)Transmission electron microscopy micrographs showing the apicoplast of intact parasites labeled with anti-GFP and 5 nm colloidal gold antibodies. *Inset* is showing a cropped and zoomed image (black box) with arrows (orange) identifying individual membranes.

This lack of processing, apicoplast localization, and the presence of three TMDs (Figure 2) suggested that UCT was likely associated with apicoplast membranes rather than imported into the organelle matrix. To test and specify UCT localization to apicoplast membranes, we labeled intact parasites expressing UCT-GFP with anti-GFP primary antibodies and colloidal gold-conjugated secondary antibodies for transmission electron microscopy analysis. Apicoplast membranes were clearly visible and decorated with gold (Figure 4C and Figure 5A-C), with UCT localizing proximal to the apparent outer membrane or appearing directly atop apicoplast membranes. This result further suggests targeting to the apicoplast membranes, and possibly the outermost membrane.

**Figure 5.**
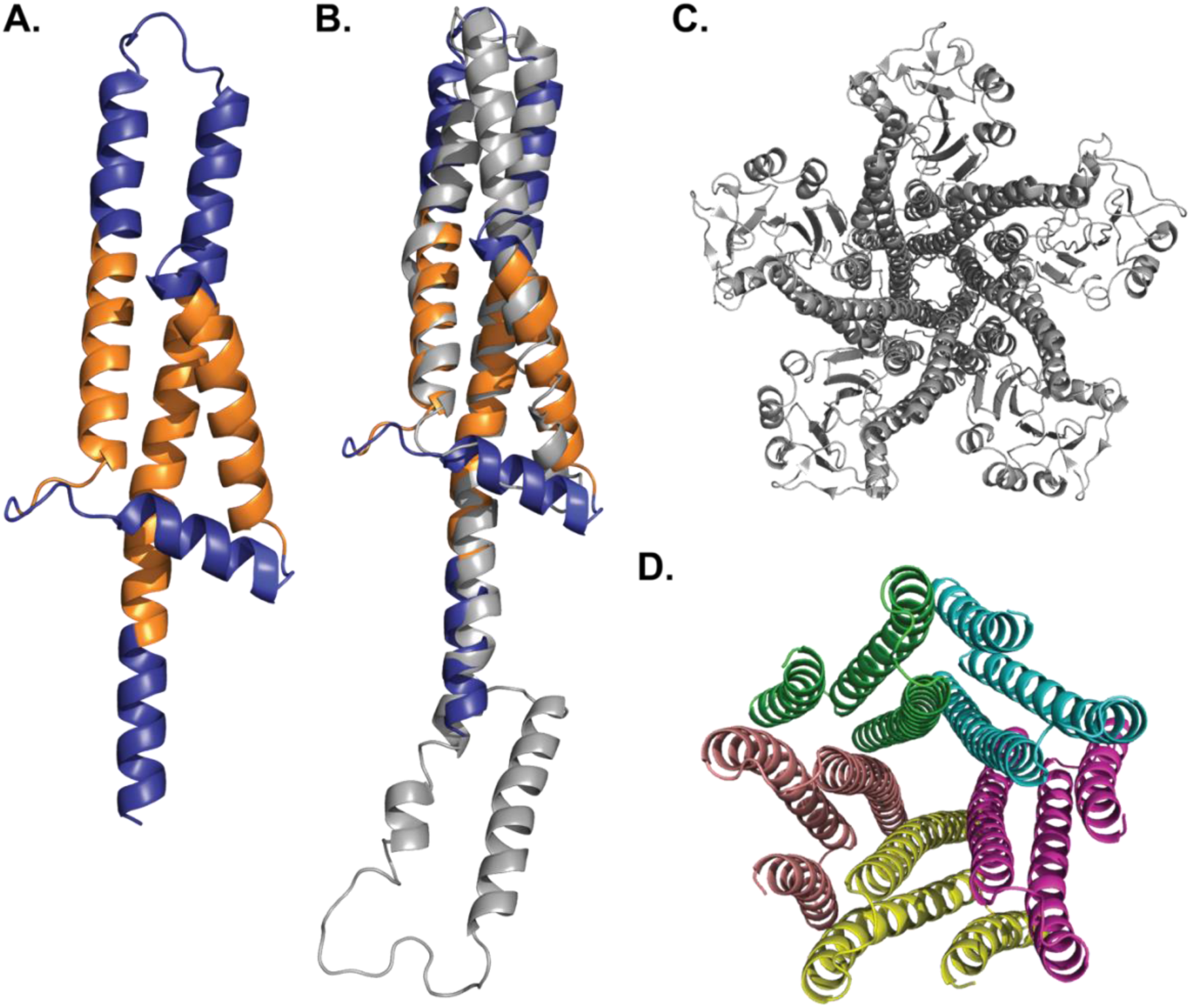
UCT has structural similarity to known metal transporters. (A) AlphaFold model of UCT with TMDs highlighted in orange. (B) Superimposition of UCT and ZntB (PDB 5N9Y, excluding N-terminal globular domain) showing similar folds, Z-score 9.1, and RMSD 3.3 Å. (C) ZntB pentameric structure. (D) Modeling of UCT as a pentamer.

### UCT has Predicted Structural Similarity to Known Metal Transporters

UCT lacks detectable sequence similarity to proteins of known function and thus lacks a predicted function. However, protein function depends more tightly on structure than on overall sequence^29^. We noted that AlphaFold predicted a 3-helical bundle for UCT, with the TMDs connected by flexible loops, which were the only sequences detected in prior mass spectrometry proteomic experiments^30–32^. To explore possible functions for UCT, we used the DALI server^33^ to perform a structural similarity search using the predicted UCT AlphaFold model^34^ (Figure 5A) as the query. This analysis identified high structural similarity between UCT and CorA-family magnesium transporter homologs from yeast^35^ (PDB 3RKG) (Figure S6A and S6E), humans^36^ (PDB 8IP4) (Figure S6B), and bacteria^37^ (PDB 5N77) (Figure S6C and S6E) and a bacterial zinc transporter^38^ (PDB 5N9Y) (Figure 5B). We also observed that UCT displayed structural similarity to the cyanobacterial ExbB monomer of the larger Ton motor complex^39^ (PDB 5SV1) (Figure S6D and S6E), which is known to transport iron. Reciprocal DALI-based searches of the *P. falciparum* structural proteome using CorA, ZntB, and ExbB structures as queries also identified substantial similarity to the AlphaFold model of UCT (Supplementary Data S2-S6). This structural similarity to metal transporters spanning multiple species suggests a possible function for UCT in apicoplast metal transport.

These transporters also form a pentameric structure with a central pore through which metal ions are shuttled. Given that the UCT monomer showed strong structural similarity to these transporters, we attempted to model a possible UCT oligomer. We observed that Alphafold modeled a pentamer only when the first 15 N-terminal amino acids were removed (Figure 5D). This observation supports the formation of a pentamer but suggests that there may be flexibility in the orientation of the N-terminal segment, which may differ in the pentameric form from what is depicted in the monomer of UCT. However, UCT shows no significant sequence homology with any of these transporters, suggesting that it may have acquired these structural features via convergent evolution of a monomeric structure favoring oligomerization.

### UCT is dispensable for the growth of asexual blood-stage parasites

We noted that the piggyBac insertional mutagenesis screen predicted a strongly negative mutant fitness score of -3.166 for UCT^40^, indicating possible blood-stage essentiality for asexual parasites. To directly test UCT function in blood-stage parasites, we edited the endogenous UCT gene locus in Dd2 parasites using CRISPR/Cas9 to encode a C-terminal 3x hemagglutinin (3HA) tag and the aptamer/TetR-DOZI system for conditional protein knockdown (KD)^41, 42^ (Figure S7A). Under this system, normal protein expression is enabled by the nontoxic small molecule anhydrotetracycline (aTc) but strongly repressed upon aTc washout. We confirmed correct genomic integration by PCR (Figure S7B). Using the 3HA tag, we performed western blot analysis to confirm endogenous UCT detection in +aTc conditions, revealing a band at ∼20 kDa, expected for full-length UCT (Figure 6A and Figure S8A). Washout of aTc resulted in a robust reduction in UCT expression, with protein levels diminished by more than 95% as determined by quantitative analysis of western blot images.

**Figure 6.**
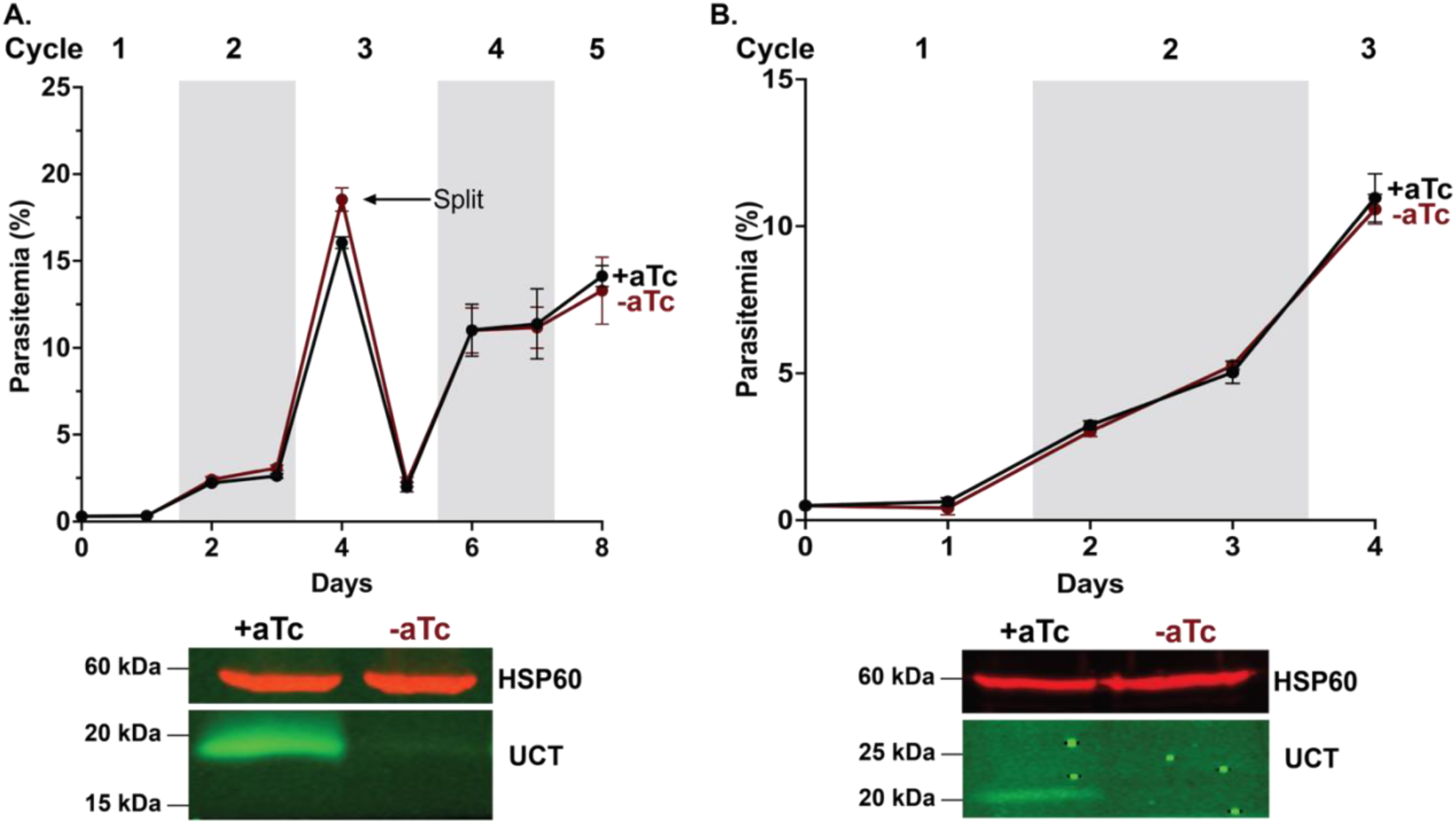
Growth of asexual blood-stage parasites does not depend tightly on UCT expression. Synchronized, continuous growth assay of UCT tagged at the endogenous locus with a 3x HA tag and the aptamer/TetR-DOZI system, and WB of tagged UCT parasites cultured in ±aTC and harvested 36 hours after synchronization and stained with anti-HA and anti-HSP60 antibodies in (A) Dd2 and (B) NF54 parasites.

In contrast to robust protein knockdown in -aTc conditions, parasites grew indistinguishably in +aTc or -aTc conditions, indicating that blood-stage growth of asexual parasites is not tightly coupled to UCT expression in Dd2 parasites (Figure 6A and Figure S7C). To further test this conclusion, we also edited NF54 parasites to enable conditional regulation of UCT. We observed a very similar result that parasites grew indistinguishably in +/-aTc conditions (Figure 6B and Figure S7B and S8B). We also tested if UCT knockdown altered parasite sensitivity to a range of drug inhibitors (fosmidomycin, doxycycline, and deferoxamine) but observed little or no shift in all cases (Figure S9).

### UCT is critical for the maturation of sexual gametocytes

Analysis of mRNA transcript levels across sexual and asexual development stages revealed that UCT transcripts are strongly upregulated in sexual gametocytes^43^, which are the parasite form that is infective to mosquitoes and thus required for transmission to the insect vector. Given this upregulation and the major metabolic remodeling that parasites undergo during development of sexual gametocytes, we hypothesized that UCT may play a critical role in gametocyte maturation. To test this hypothesis, we edited the endogenous UCT gene locus in inducible NF54/iGP2 parasites^44^ to encode the same conditional protein KD system used in Dd2 and NF54 parasites. We confirmed correct genomic integration by PCR (Figure S10).

Gametocyte induction in the NF54/iGP2 system is under the control of the *glmS* riboswitch, such that glucosamine washout causes ∼75% of asexual parasites to commit to sexual development. Synchronized ring-stage parasites were induced by glucosamine washout, split into +/- aTc conditions, and allowed to expand. To monitor gametocyte maturation, we killed off asexual parasites by treatment during days 1-6 with N-acetyl-D-glucosamine (GlcNac) (Figure 7A). We performed western blot analysis using the 3HA tag to confirm endogenous UCT detection in +aTc conditions and reduced expression in -aTc conditions (Figure 7B and Figure S11). In contrast to our observations in asexual parasites, we observed a substantial defect in gametocyte maturation upon UCT knockdown (Figure 7). This defect included a strongly decreased percentage of stage IV and V gametocytes and an overall increase in gametocytes with irregular morphology, without significant difference in stage III gametocyte formation. (Figure 7B and Figure S12-14). These results suggest that UCT expression and function are critical for the full development of sexual gametocytes beyond stage III to mature stage V parasites required for transmission to the mosquito vector.

**Figure 7.**
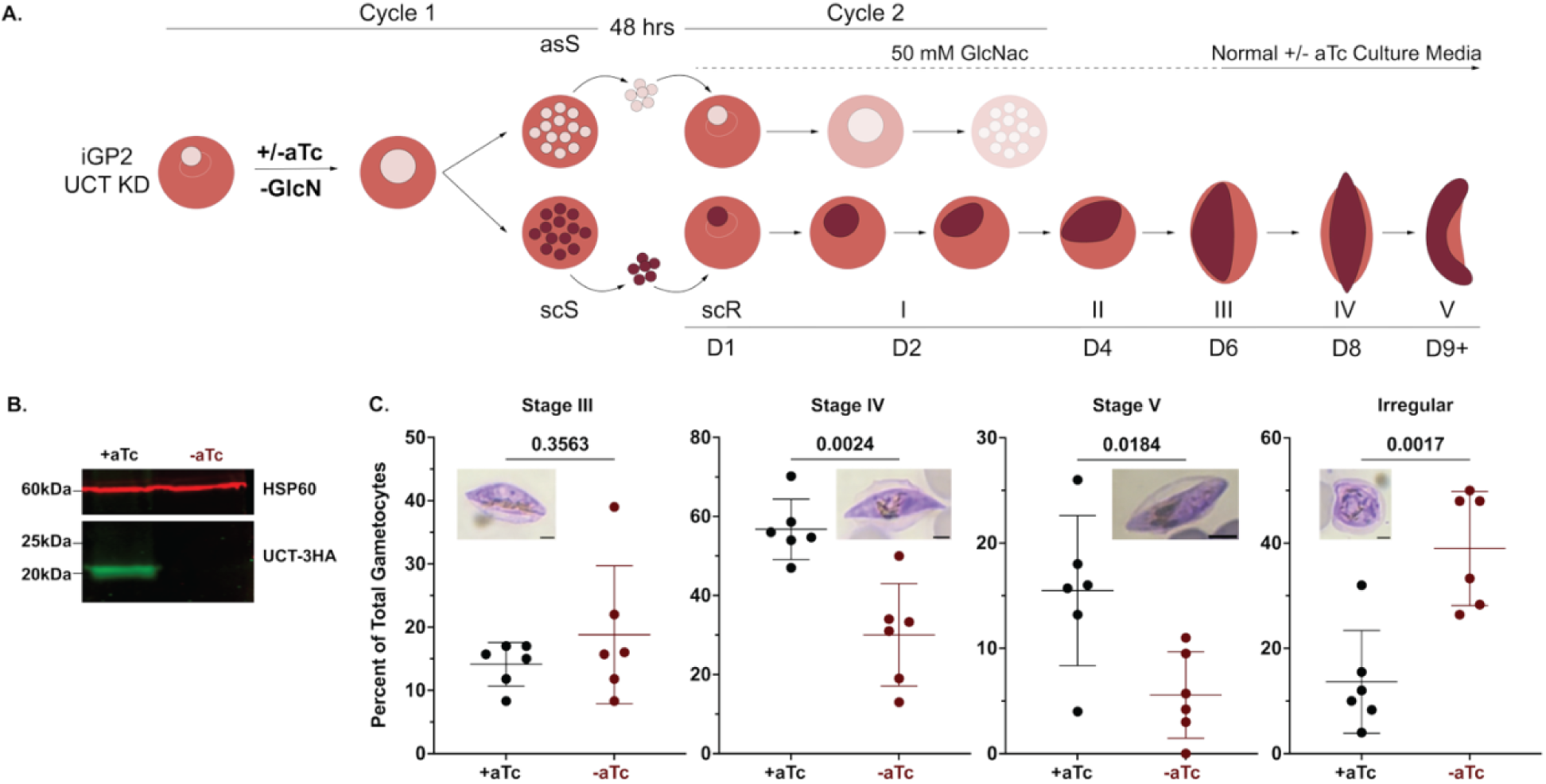
UCT is necessary for gametocyte maturation. (A) Schematic depiction of the culture protocol used to induce gametocytes using glucosamine washout (-GlcN) in +/-aTc conditions. After 48 hours, asexual stage schizonts (asS) and sexually committed schizonts (scS) rupture and reinvade, and sexually committed rings (scR) mature over 9+ (D1-9+) days into gametocytes. Starting at 48-hours (D1), 50 mM N-acetyl-D-glucosamine (GlcNac) was added for 6 days to arrest asexual parasite progression. Asexual parasites are depicted in light pink, sexually committed parasites are depicted in burgundy, and shaded graphics depict blocked growth of asexual parasites. (B) Western blot of tagged UCT parasites cultured in ±aTC, induced with GlcN, harvested after 9 days, and stained with anti-HA and anti-HSP60 antibodies. (C) Gametocyte stages were quantified by Giemsa-stained blood smears of three independent clones on day 9 after growth in +/- aTc conditions by blinded analysis of ≥70 gametocytes per clonal condition. Scale bar = 2.5 microns. P values determined by t-test are given in the above data sets.

## Discussion

Since the discovery of the apicoplast, extensive work has been done to define its key metabolic functions and its essentiality for parasite viability. Nevertheless, the transporters that mediate nutrient movement across four membranes to support biochemical pathways in the organelle remain sparsely defined and represent a key gap in our understanding of this compartment^9^. Here, we identify a previously uncharacterized protein with structural similarity to known metal transporters in other organisms. This finding provides a promising candidate for understanding metal-import function into the apicoplast. It also demonstrates the power of structure-based bioinformatic approaches to suggest functions for highly divergent parasite proteins that defy functional annotation by sequence analysis alone. Our discovery provides a biochemical foothold for both testing specific metal-transport properties of UCT and for unveiling broader features of metal and nutrient transport into the apicoplast.

### UCT as a putative apicoplast target of dox

We identified UCT as the top protein interactor with tetracycline affinity resin in two independent pull-downs using *P. falciparum* cell lysates. This protein was selectively enriched in the absence of dox, eluted with free dox, and depleted upon pretreatment of lysates with free dox before affinity isolation, suggesting a specific interaction with dox. A similar approach was used to identify tetracycline interactions with 80S ribosomes in human cells^19^, but this tool has not been used previously in malaria parasites. Tetracyclines like dox have been assumed to target apicoplast ribosomes exclusively in *Plasmodium*, but we did not identify ribosomal proteins as strongly enriched in our affinity isolations. This observation may suggest that such ribosomal interactions in parasites are lower affinity, are labile to disruption under non-equilibrium pull-down conditions, and/or are disrupted by conjugation of the tetracycline ligand to an agarose resin despite a flexible linker. Our prior work identified apicoplast-specific activity by dox beyond the expected inhibition of apicoplast ribosomes^15^. Identification of UCT as a tetracycline-interacting protein suggests that dox may inhibit UCT function as an additional apicoplast target. However, the dispensable function of UCT for blood-stage parasites strongly suggests that UCT is unlikely to be the exclusive or dominant target of the higher 10 µM dox dose that leads to first-cycle parasite death.

### Trafficking and localization of UCT to the apicoplast

We confirmed that UCT is targeted to the apicoplast, where it localizes closely with apicoplast ACP. Most proteins that function within the apicoplast contain N-terminal signal and transit peptides that are proteolytically cleaved upon import into the endoplasmic reticulum and the apicoplast^23, 24^, respectively. UCT lacks a canonical signal peptide, but its N-terminus contains multiple positively charged Lys and Arg residues following the first transmembrane domain, as expected for a transit peptide. We observed that the N-terminal 42 residues are sufficient to confer apicoplast targeting, suggesting that the N-terminal transmembrane domain can function as an effective signal peptide.

In contrast to most apicoplast proteins, neither full-length UCT nor the N-terminal 42 residues fused to GFP showed evidence of proteolytic processing upon apicoplast targeting. This observation, the presence of multiple transmembrane domains, and immunogold electron microscopy that revealed targeting to apicoplast membranes collectively suggest that UCT localizes to one of the outer membranes with the N-terminus not exposed to the matrix, where proteolytic processing occurs. In this regard, UCT is similar to the outer triosphosphate transporter (oTPT), which localizes to the outermost apicoplast membrane without proteolytic processing^28^. We posit that UCT may also localize to the outer membrane, but further testing is required to confirm this model.

In contrast to oTPT, which lacks obvious apicoplast-targeting features, the N-terminus of UCT is sufficient for apicoplast targeting without processing. UCT appears to transit the ER without involving the Golgi. Its localization is independent of brefeldin A treatment (Figure S15), and apicoplast disruption results in dispersed UCT localization in cytoplasmic foci similar to apicoplast ACP. We propose that the first N-terminal transmembrane domain of UCT functions as a de facto signal peptide for ER entry. Its lack of processing in the ER may be due to the slightly recessed position of this TMD relative to the N-terminus and the presence of bracketing basic residues at either end of this hydrophobic segment that prevent cleavage by the ER signal peptidase. We have ongoing experiments to further test and dissect these features. Finally, we note that the lack of proteolytic processing is consistent with our affinity-isolation of UCT from lysates of parasites with both intact and disrupted apicoplasts.

### UCT functions in the apicoplast

UCT is highly conserved across *Plasmodium* species, but sequence alignments and BLAST analyses provided little insight into its function. However, UCT retains structural similarity to oligomeric CorA-family magnesium and zinc transporters and the cyanobacterial ExbB iron transporter^35–39^. Structural similarity to each transporter includes the TMDs of UCT but corresponds to regions of the soluble domains of CorA, ZntB, and ExbB. This 3-helical structural similarity between UCT and CorA/ExbB transporters, despite the lack of detectable sequence similarity, may reflect convergent evolution of a structural motif that favors oligomerization with a central pore. Such a pentameric assembly is a key feature of CorA/ExbB transporters and is supported by UCT modeling. Additionally, CorA and ZntB transporters have N-terminal globular domains absent in UCT, which may suggest that UCT function does not require this additional domain or that UCT has additional interacting proteins at the apicoplast that contribute to its function.

Preliminary AlphaFold modeling indicates that UCT assembles into a pentameric structure when the N-terminal region is partially removed (Figure 5). However, our data suggest that UCT is not N-terminally processed, which may indicate N-terminal flexibility beyond that modeled by AlphaFold in its monomeric state and/or a conformational shift upon oligomerization with an interaction partner. This structural scenario has previously been observed with gasdermin D, which contains flexible loops that undergo a conformational change during the transition from prepore to pore formation during the release of inflammatory cytokines^45^. We have ongoing work to purify UCT under native conditions for negative-stain electron microscopy to assess its oligomeric state. It remains a critical challenge to test the metal-transport properties of UCT directly and to identify key interactors that may be critical for UCT function.

Direct knockdown of UCT had no detectable impact on the growth or viability of asexual blood-stage parasites, in contrast to insertional mutagenesis studies in *P. falciparum* that suggested a negative fitness score for UCT disruption^40^. Based on the model of UCT function in metal transport, this result may indicate functional redundancy in metal-uptake pathways into the apicoplast during blood-stage infection. Indeed, distinct metal transporters can exhibit promiscuity in their metal substrates, and parasites may have evolved redundant mechanisms to import key nutritional metals into the apicoplast. A central example of promiscuous metal transport is provided by eukaryotic DMT1, which primarily transports ferrous iron but has some affinity to transport other divalent metals (e.g., Zn^2+^, Mn^2+^, Co^2+^, etc.)^46, 47^. This promiscuity is also a feature of the CorA family of transporters, which can transport additional divalent metals (e.g., Co^2+^, Ni^2+^, and Cd^2+^) in addition to Mg^2+^ and Zn^248, 49^. Another possibility is that low levels of UCT expression are sufficient to maintain parasite viability despite stringent knockdown, as is the case for the essential endoplasmic reticulum protease, plasmepsin V^50–52^.

UCT expression is upregulated 10-20-fold in stage V sexual gametocytes and in mosquito stages of the parasite life cycle^43^, suggesting a more crucial function in these stages. We observed that UCT knockdown in gametocytes resulted in a substantial maturation defect, with fewer gametocytes progressing to stage IV and V and a significant increase in irregular gametocytes. These data suggest an essential role for UCT in the development of sexual blood-stage gametocytes. The biochemical origin of this defect upon UCT knockdown remains to be understood. Given UCT localization to the apicoplast and its proposed role in iron/metal import, we hypothesize that defective gametocyte development reflects apicoplast dysfunction and impaired isoprenoid synthesis. We noted that hemozoin crystals in UCT knockdown gametocytes often appeared to adopt distinct morphologies compared to healthy gametocytes, which might suggest defects in food vacuole function due to impaired isoprenoid synthesis in the apicoplast, as described for asexual parasites^53, 54^. We have ongoing experiments to test UCT function during parasite growth within mosquitoes.

### Closing perspective

Identification of apicoplast membrane transporters hidden among unannotated proteins in the parasite genome remains a key challenge for fully understanding nutrient-uptake mechanisms in this organelle. Recent efforts have used sequence- and structural-similarity searches to identify candidate apicoplast transporters, but few (only 5 of 28) have been experimentally investigated^9^. This current list of putative transporters is likely insufficient to fully account for metabolite fluxes mediated by the apicoplast transportome. UCT has not previously been identified as a candidate apicoplast transporter. We propose that UCT functions as an apicoplast metal transporter, possibly for the uptake of iron and/or other metals. Its heightened expression and functional contributions in sexual gametocytes and mosquito stages suggest a critical role in parasite transmission to the insect vector and new human hosts. We also highlight the crucial role of structure-based similarity searches in suggesting a functional role for UCT, and we expect these approaches to advance understanding of other *P. falciparum* proteins that lack functional annotation based solely on sequence analysis. With the continuing development of antimalarial drug resistance, it will remain critical to identify new functions for apicoplast and other parasite proteins that can be exploited as novel drug targets for antimalarial therapies.

## Materials and Methods

### Parasite culturing and transfections

*Plasmodium falciparum* Dd2^55^, PfMev^26^, NF54^56^, or NF54/iGP2^44^ parasites were maintained at 2% hematocrit and cultured in human erythrocytes acquired from the University of Utah Hospital blood bank (Salt Lake City, UT) at 37°C in either 90% N_2_/5% CO_2_/5%O_2_ or 5% CO_2_ balanced with ambient air. All culturing was performed in Roswell Park Memorial Institute medium (RPMI-1640, Thermo Fisher 23400021), 2.5g/L Albumax 1 Lipid-Rich BSA (Thermo Fisher 11020039), 15 mg/L hypoxanthine (Sigma H9636), 110 mg/L Sodium pyruvate (Sigma P5280), 1.19g/L HEPES (Sigma H4034), 2.52 g/L sodium bicarbonate (Sigma S5761), 2 g/L glucose (Sigma G7021), and 10 mg/L gentamicin (Invitrogen Life Technologies 15750060), as described previously^15, 57, 58^. Uninfected erythrocytes were transfected with 1x cytomix containing 50-150 μg of midi-prepped and ethanol-precipitated DNA via electroporation in 0.2 cm cuvettes using a Bio-Rad Gene Pulser Xcell system (0.31 kV, 925 μF). Parasite-infected erythrocytes were then added, allowed to rupture and infect transfected erythrocytes for 48 hours, and then subjected to drug pressure to positively select for plasmid uptake. Selectable markers were encoded on the transfected plasmid and included human DHFR^59^ or blasticidin-S deaminase (BSD)^60^. Cultures were maintained in 5 nM WR99210 and 6 μM BSD, respectively.

Parasites that contained UCT (PF3D7_1248300) tagged with the aptamer/TetR-DOZI cassette^41^ were cultured in 1 μM anhydrotetracycline (aTc, Caymen chemicals 10009542). All genetically modified parasite lines were genotyped by PCR. For apicoplast drug-induced disruption experiments, parasites were cultured for ≥4 days in the presence of 10 μM doxycycline (Sigma, D9891) and 200 μM isopentenyl pyrophosphate (Isoprenoids, IPP001) for Dd2 or 50 μM mevalonate (Cayman Chemicals, 20348) for PfMev parasites.

### Tetracycline Affinity Pulldown

Two 75 mL cultures of untreated Dd2 and one culture of apicoplast-disrupted Dd2 parasites were harvested at ∼15% parasitemia by centrifugation, incubated with 0.05% saponin (Sigma S7900) in 1x phosphate buffered saline (PBS) for 5 minutes at room temperature. Parasites were then lysed in 1% v/v Triton X-100 with protease inhibitors (Invitrogen) by sonication (3 x 10 pulses, 50% duty cycle, 50% power) on a Branson microtip sonicator. Sonicated samples were then incubated for 2 hours at 4°C with continuous rotation, followed by centrifugation (17,000 x g, 30 minutes). One sample of Dd2 parasites was treated with 200 μM doxycycline during the lysis stage. Total protein concentration in lysates was determined by Lowry assay. Clarified lysates containing 500 μg of total protein were mixed with 50 μL equilibrated immobilized tetracycline affinity resin (G-Biosciences, 786-1331) and incubated at 4°C for 2 hours with continuous rotation. Immobilized tetracycline agarose was pelleted by centrifugation, the supernatant was removed by aspiration, and the agarose was washed 3 times in 1% Triton X-100/1x PBS in autoclaved, ethanol-treated Eppendorf tubes. Each wash was performed in a new tube. Bound proteins were then eluted with 100 μL of 100 μM doxycycline for one hour at 37°C. Proteins were then precipitated using 100% trichloroacetic acid (Sigma 76039) to a final concentration of 20%, incubated on ice for 1 hour in Eppendorf tubes. Tubes were then centrifuged at 13,000 RPM for 25 minutes at 4°C. The supernatants were removed by aspiration, and protein pellets were washed once in 500 μL of cold acetone. Pellets were subsequently air-dried for 30 minutes.

### Mass Spectrometry

Protein samples were reduced and alkylated using 5 mM Tris (2-carboxyethyl) phosphine and 10 mM iodoacetamide, respectively, and then enzymatically digested by sequential addition of trypsin and lys-C proteases, as previously described^61^. The digested peptides were desalted using Pierce C18 tips (Thermo Fisher Scientific), dried, and resuspended in 5% formic acid. Approximately 1 μg of digested peptides was loaded onto a 25-cm-long, 75-μm inner diameter fused silica capillary packed in-house with bulk ReproSil-Pur 120 C18-AQ particles, as described previously^62^. The 140-min water-acetonitrile gradient was delivered using a Dionex Ultimate 3,000 ultra-high performance liquid chromatography system (Thermo Fisher Scientific) at a flow rate of 200 nL/min (Buffer A: water with 3% DMSO and 0.1% formic acid, and Buffer B: acetonitrile with 3% DMSO and 0.1% formic acid). Eluted peptides were ionized by the application of distal 2.2 kV and introduced into the Orbitrap Fusion Lumos mass spectrometer (Thermo Fisher Scientific) and analyzed by tandem mass spectrometry. Data were acquired using a Data-Dependent Acquisition method consisting of a full MS1 scan (resolution = 120,000) followed by sequential MS2 scans (resolution = 15,000) for the remainder of the 3-s cycle time. Mass spectrometry data were processed using the MaxQuant bioinformatics pipeline^63^ , with peptide identification carried out by the Andromeda search engine against the *Plasmodium falciparum 3D7* reference proteome (UniProt ID: UP000001450). Key settings included: a maximum of two missed cleavages, a false discovery rate (FDR) of 1% for both peptides and proteins, label-free quantification (LFQ) enabled with a minimum LFQ ratio count of 1, and parent and precursor ion tolerances of 20 and 4.5 ppm, respectively. The proteomics data are deposited in the MassIVE data repository (https://massive.ucsd.edu) under the identifier MSV000101739. Protein enrichment values were calculated as the log_2_ ratio of spectral counts or intensity values for the indicated protein in the experimental versus negative-control datasets. Protein spectral count values of zero were arbitrarily converted to one for purposes of calculating a log_2_ enrichment ratio.

### Cloning

The gene encoding UCT (PF3D7_1248300) was PCR amplified from Dd2 *P. falciparum* parasites using primers with 20 base pair overhangs for ligation-independent insertion into the XhoI and AvrII sites of pTEOE containing the human DHFR selection cassette^64^ and in frame with a C-terminal GFP or mScarlet marker. Primer sequences are provided in Table S1. All primers to enable episomal expression were designed with 20-base pair (bp) overhangs into the corresponding expression plasmid. CRISPR/Cas9 was used to tag the endogenous UCT gene in Dd2 or NF54 parasites to encode a 3x HA C-terminal tag and the 10x aptamer/TetR-DOZI conditional regulation system via repair by double-crossover homologous recombination^41, 42^. The plasmid was assembled in three total fragments and subsequently combined via assembly PCR. Briefly, the first fragment (370 bp) contained homologous overlap to the 3’ untranslated region (UTR) of UCT. Coding sequence (CDS) fragment 1 (254 bp) and CDS fragment 2 (196 bp) were designed using primers to introduce shield mutations at the gRNA binding region upstream of the PAM sites for CDS gRNA 2 and 3. All primers were designed to include a 20 bp overlap with the expression plasmid or the corresponding fragment for assembly PCR. A pAIO plasmid with each of 3 gRNA sequences located either in the UCT CDS or in the 3’UTR was cloned, and plasmids were mixed during transfection (CDSgRNA2:AGTGCGAATGAGCTAATCAC) (CDSgRNA3:TAGTGCGAATGAGCTAATCA)(UTRgRNA5:CAAATGATAAAATGTGTAG C). All cloning reaction mixes were transformed into chemically competent Top10 cells, and clones were selected based on carbenicillin resistance (Gold Biotechnology C-103-1). For recombinant expression in *E. coli*, UCT-GFP was amplified from plasmid DNA of UCT-GFP/pTEOE for insertion between the NdeI/XhoI sites of pET24a. Due to the presence of an NdeI restriction site in the GFP cassette sequence, cloning from plasmid DNA was designed to exchange the NdeI site to a XhoI site. Correct plasmid sequences were confirmed by Sanger sequencing or whole plasmid sequencing performed by Plasmidsuarus using Oxford Nanopore Technology.

### Parasite growth assays

Parasite continuous growth assays were performed using sorbitol-synchronized parasites, seeded to ∼0.5% parasitemia, and allowed to expand over multiple days with daily media changes. Culture parasitemia was determined daily via flow cytometry by diluting 20 μL from individual parasite culture wells of each biological triplicate into 180 μL of 1.0 μg/mL acridine orange (Invitrogen Life Technologies A3568) in PBS. Analysis was performed on a BD FACSCelesta system by monitoring FSC-A, SSC-A, PE-A, FITC-A, and PerCP-Cy5-5-A channels. Daily measurements were plotted as a function of time and graphed using GraphPad Prism 10. For EC_50_ determinations, sorbitol synchronized parasites were seeded at 0.05% parasitemia and incubated with variable drug concentrations for 48-hours without media changes. Parasitemia was measured as described above and plotted as a function of the log of drug concentration (μM) and fit to a four-parameter dose-response curve using GraphPad Prism 10.

### Gametocyte Maturation Assays

Gametocyte maturation assays were performed as described previously^44^. Briefly, gametocyte maturation assays were performed using synchronous NF54/iGP2 ring-stage cultures at 5-7% parasitemia, washed in normal culture media to remove GlcN and wash out aTc from the media. Cultures were then split into +/-aTc conditions. 48 hours post-synchronization, on day 1 of gametocytogenesis, 50 mM N-acetyl-D-glucosamine (MP Biomedicals 194604) was added to ring-stage cultures and changed daily for 6 consecutive days to eliminate remaining asexual parasites. Starting day 7, parasites were cultured in normal +/- aTc culture media. Gametocyte maturation was assessed on day 9 in three independent clonal UCT KD lines by blinded visual inspection of ≥70 gametocytes per clonal condition of Giemsa-stained blood smears using an EVOS M5000 microscope.

### SDS-PAGE and western blot analysis

Asynchronous parasite cultures were grown to between 10-15% parasitemia in 50-100 mL cultures and harvested by centrifugation followed by 0.05% saponin in PBS for 5 minutes at room temperature and pelleted by centrifugation at 4,000 RPM for 30 minutes at 4°C. Samples were lysed using sonication (10 pulses twice, 50% duty cycle, 25-30% power) on a Branson microtip sonicator in PBS lysis buffer (1% v/v Triton x-100 and/or 0.1% SDS) in the presence of protease inhibitors (Invitrogen A32955 or Selleckchem B14001). Lysate was then incubated at 4°C for 1-2 hours. Recombinantly expressed protein was obtained by induction of BL21 (DE3) *E.coli* grown in LB medium to an OD of 0.3-0.6 with 1 mM IPTG at 20°C overnight. The cell pellet was harvested by centrifugation (4,500 RPM, 10 min) and lysed as described above. Lysates were clarified by centrifugation (17,000 rcf x g, 30 minutes), mixed with SDS sample buffer, heated at 65°C for 5 minutes and then separated on an SDS-PAGE gel (10%, 12%, 15%, or 4-16% gradient). Proteins were transferred onto a nitrocellulose membrane using a wet transfer for 1 hour at 100 V. Membranes were blocked in 5% w/v non-fat milk in PBS. Membranes were stained with primary antibodies: goat anti-GFP (antibodies.com LLC A121560), rat anti-HA (Sigma 11867423001), and rabbit anti-HSP60 (Novus Biologicals NBP2-12734) diluted to 1:2,000 in 1% w/v non-fat milk in PBS with 0.1% Tween 20 (Sigma P1379). Membranes were subsequently washed thrice in PBS with 0.1% Tween 20. Membranes were then stained in secondary antibodies: donkey anti-goat (LICOR 926-32214), donkey anti rat (Thermo Fisher SA5-10032) and goat anti-rabbit (LICRO 925-68071) or donkey anti-rabbit (LICOR 926-68023) diluted to 1:10,000 in 1% w/v non-fat milk in PBS with 0.1% Tween 20 (Sigma P1379).

### Microscopy

For all untreated live-cell experiments, asynchronous parasites were analyzed. For all doxycycline- and Mev/IPP-treated live-cell experiments, parasite samples were collected 36 hours post-synchronization with 5% D-sorbitol (Sigma S7900). For brefeldin A sensitivity experiments, parasite samples were collected 18 hours post-synchronization and treatment with 5 µg/mL brefeldin A (Sigma B7651). Parasite nuclei were visualized via incubation with 1–2 µg/mL Hoechst 33342 (Thermo Scientific Pierce 62249) for 10 minutes at room temperature. UCT was visualized in Dd2^55^ parasites via a C-terminal GFP tag and in PfMev^26^ via a C-terminal mScarlet tag. The apicoplast in PfMev parasites was visualized using ACP_L_-GFP, expressed as an organelle marker in these parasites.

For all IFA imaging, the parasite apicoplast was visualized via a custom polyclonal rabbit anti-ACP antibody^65^ diluted to 1:50 and goat anti-rabbit fluorescent secondary antibody (Invitrogen R37117) diluted to 1:1,000, and the nucleus was stained with ProLong Gold Antifade Mountant with DAPI (Invitrogen Life Technologies P36931). UCT-GFP and UCT_N-Term_-GFP were visualized with a goat anti-GFP (antibodies.com LLC A121560) diluted to 1:250 and donkey anti-goat (Abcam AB150129) diluted to 1:1,000 fluorescent secondary antibody. Images were taken on DIC/bright field, DAPI, GFP, and RFP channels using either an EVOS M5000 or Spinning Disk Confocal imaging system. Fiji/ImageJ was used to analyze and process all images. Intensity plots were generated using the “plot profile” function using a shared region of interest (white line) on each channel. All image adjustments were made on a linear scale, including brightness and contrast.

For immunogold transmission electron microscopy, 40 ml of parasite UCT-GFP samples were prepared by passage over a magnet column (Miltenyi Biotech, 130-042-901), pelleted by centrifugation at 2,000 RPM for 3 minutes and fixed for 2 hours at room temperature in 2% glutaraldehyde and 1% paraformaldehyde (85 mM HEPES, 25 mM PIPES, pH 6.9, 370 milliosmole). After fixation, the cells were washed to remove residual fixative and pelleted by centrifugation. Instead of gelatin embedding, the cells were encapsulated in ethylene-vinyl alcohol (EVOH) copolymer hollow fibers^66^ and infused with 15% polyvinylpyrrolidone (PVP 10,000) and 1.7 M sucrose. Segments of the EVOH fibers containing highly concentrated, cryoprotected cells were mounted on specimen stubs and cryo-sectioned at −80 °C using a Leica UC7 ultracryomicrotome. Ultrathin cryosections (70 nm) were immunolabeled following established procedures^67^. The primary antibodies and gold-conjugated markers used were rabbit anti-ACP (1:1000) detected with 10 nm protein-A gold (1:25; UMC, The Netherlands), and anti-GFP (1:100; Abcam 5450) detected with 5 nm gold-conjugated anti-goat antibody (1:30; Cytodiagnostics Inc.). Transmission electron microscopy (TEM) images were acquired on a JEOL JEM-1400Plus operated at 120 kV.

### Structural Similarity Searches and Modeling

The predicted monomeric structure of UCT was obtained from the AlphaFold Protein Structure Database^34^ , and experimentally determined structures of CorA magnesium, ZntB zinc, and ExbB iron transporters were obtained from RCSB.org^68^. All structural similarity data were obtained from the DALI server^33^. Protein models were visualized and analyzed with The PyMOL Molecular Graphics System, Version 3.1.6.1, Schrödinger LLC.

## Acknowledgements

We thank Till Voss (Swill Tropical and Public Health Institute) for providing NF54/iGP2 parasites, Radoslaw Omelianczyk for counting gametocytes, and members of the Sigala lab for helpful discussions. This work was supported by NIH grants R21AI185746 (to PAS) and R35GM153408 (to JAW). SN was supported by a diversity supplement to R35GM133764 (to PAS) and by an NSF graduate research fellowship (DGE-1656518). DNA synthesis and Sanger sequencing, fluorescence and electron microscopy, and flow cytometry were performed using core facilities at the University of Utah.

## Notes

ftp://massive-ftp.ucsd.edu/v12/MSV000101739/

### Competing Interest Statement

The authors have declared no competing interest.

