## Supporting Information for "Identification of a Putative Metal Transporter in the Apicoplast of Malaria Parasites"

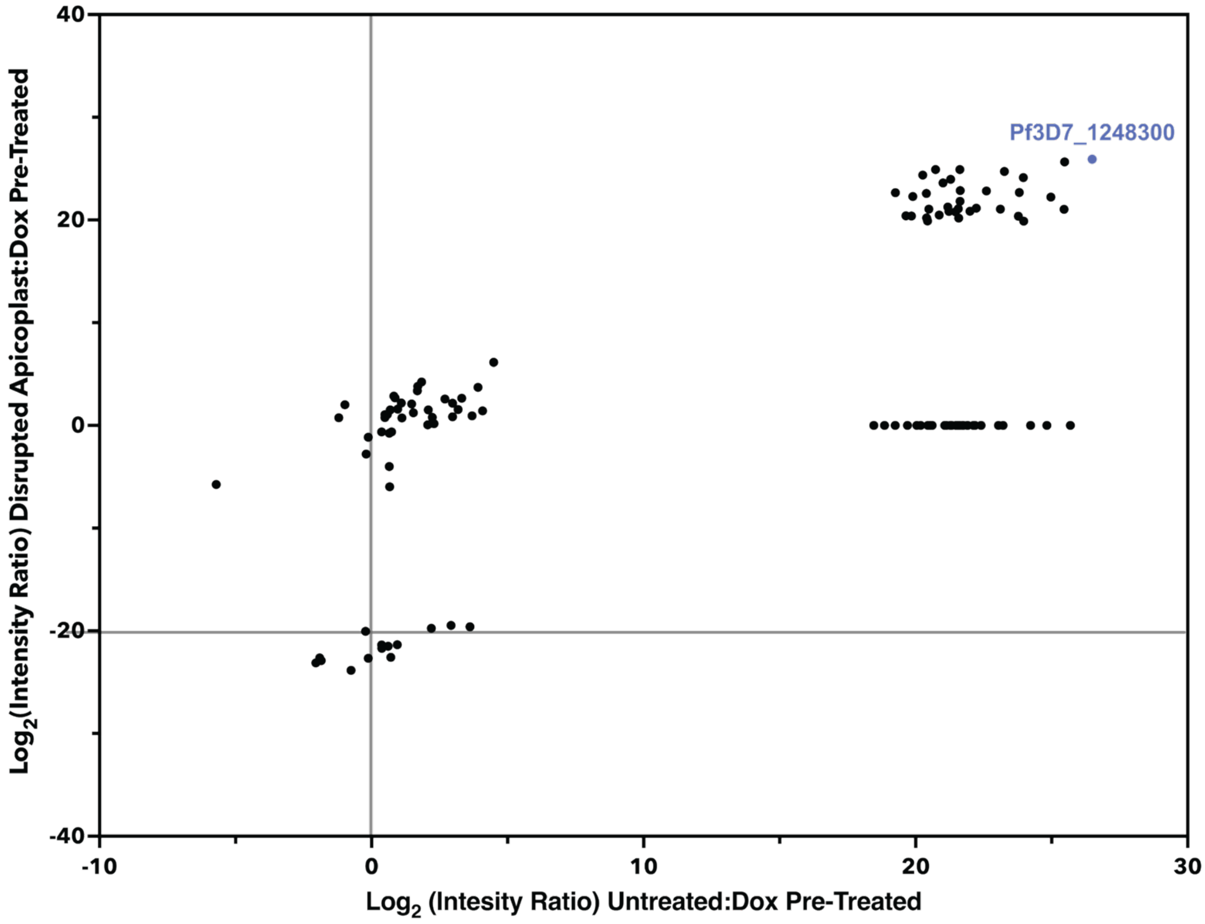


**Figure S1.** Tetracycline affinity pull-down data set. Proteomic results showing the $log_2$ ratio of spectral abundance for proteins identified in untreated versus dox-pretreated or disrupted apicoplast versus dox-pretreated lysates, including the entire protein hit dataset. Pf3D7_1248300 (UCT) highlighted in blue.

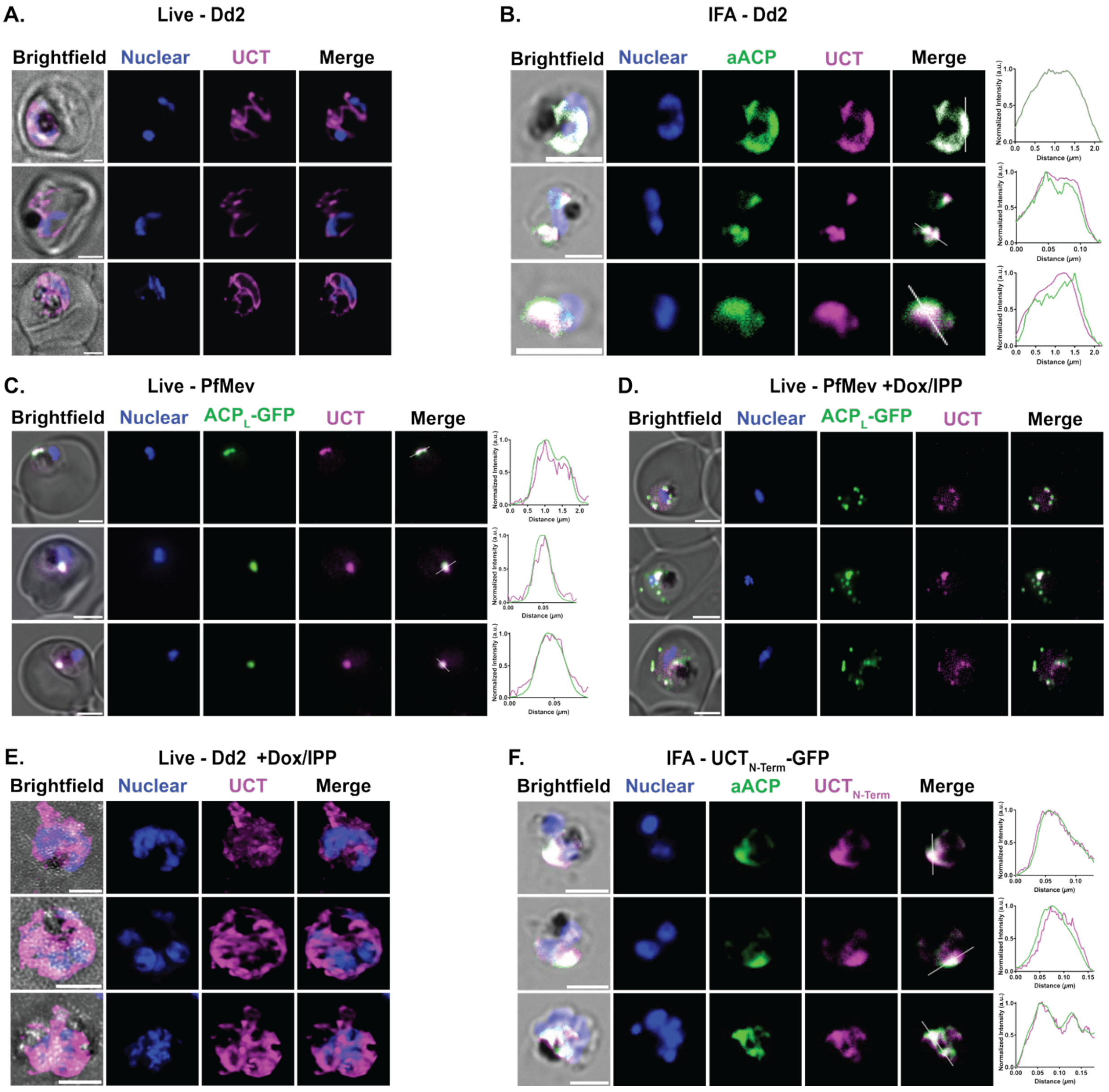


**Figure S2.** UCT localizes to the apicoplast. (A) Live-parasite fluorescent microscopy of UCT-GFP showing expected organelle branching (B). Immunofluorescent microscopy images of UCT-GFP showing co-localization with apicoplast ACP (aACP) (C). Live-parasite fluorescent microscopy of UCT-mScarlet (mS) showing co-localization with ACP$_L$-GFP. (D) Live-parasite fluorescent microscopy of UCT-mS in PfMev parasites treated with doxycycline and mevalonate showing dispersed foci of UCT-mS and ACP$_L$-GFP upon apicoplast disruption. (E) Live-parasite fluorescent microscopy of UCT-GFP treated with doxycycline and IPP showing dispersed foci of UCT upon apicoplast disruption. (F) Immunofluorescent microscopy images UCT$_{N\text{-term}}$-GFP showing co-localization with aACP. Scale bar = 2.5 microns.

A.

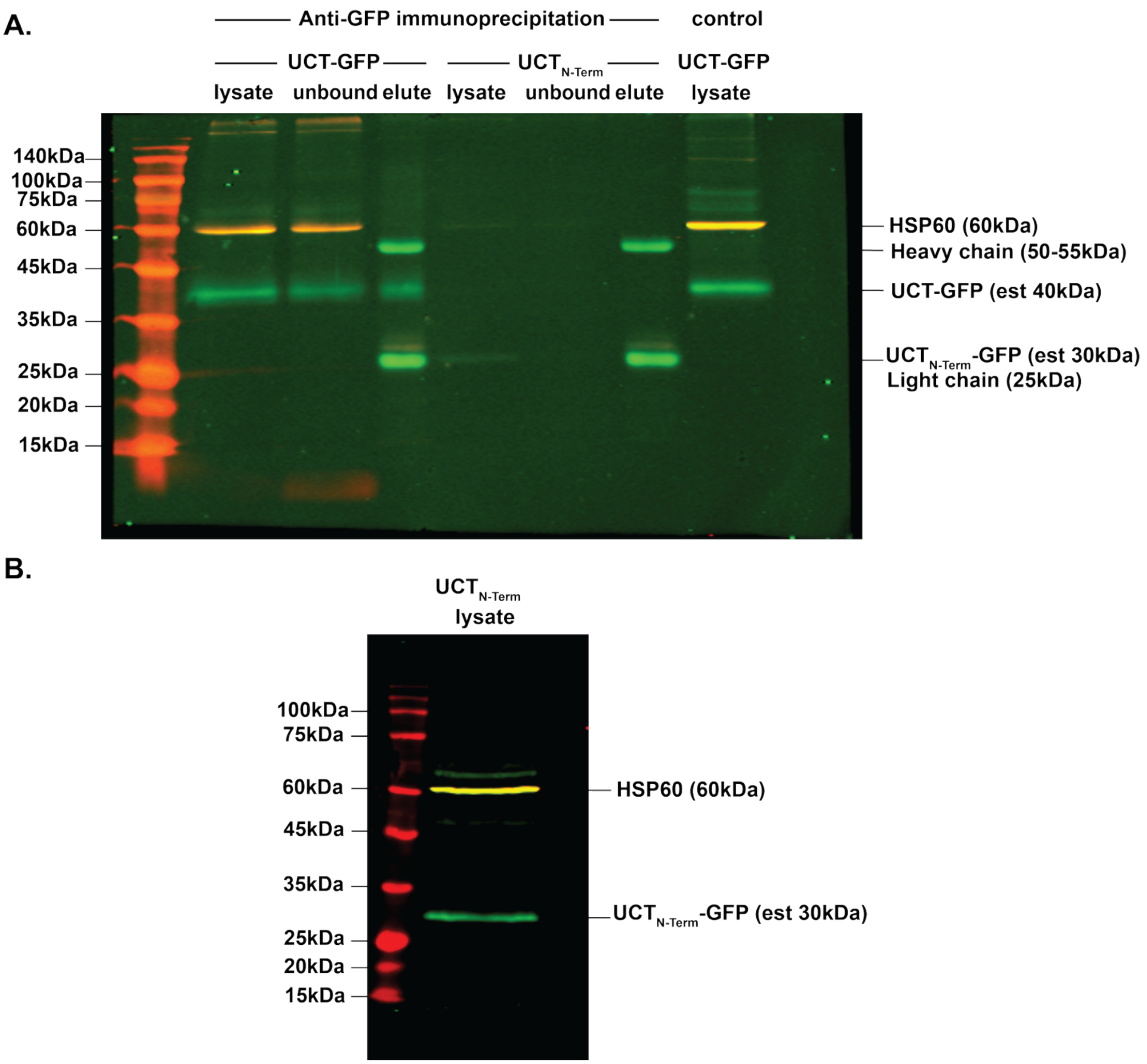


**Figure S3.** UCT western blot analysis of (A) full-length UCT-GFP via an anti-GFP immunoprecipitation compared to UCT$_{N-Term}$-GFP and (B) UCT$_{N-Term}$-GFP.

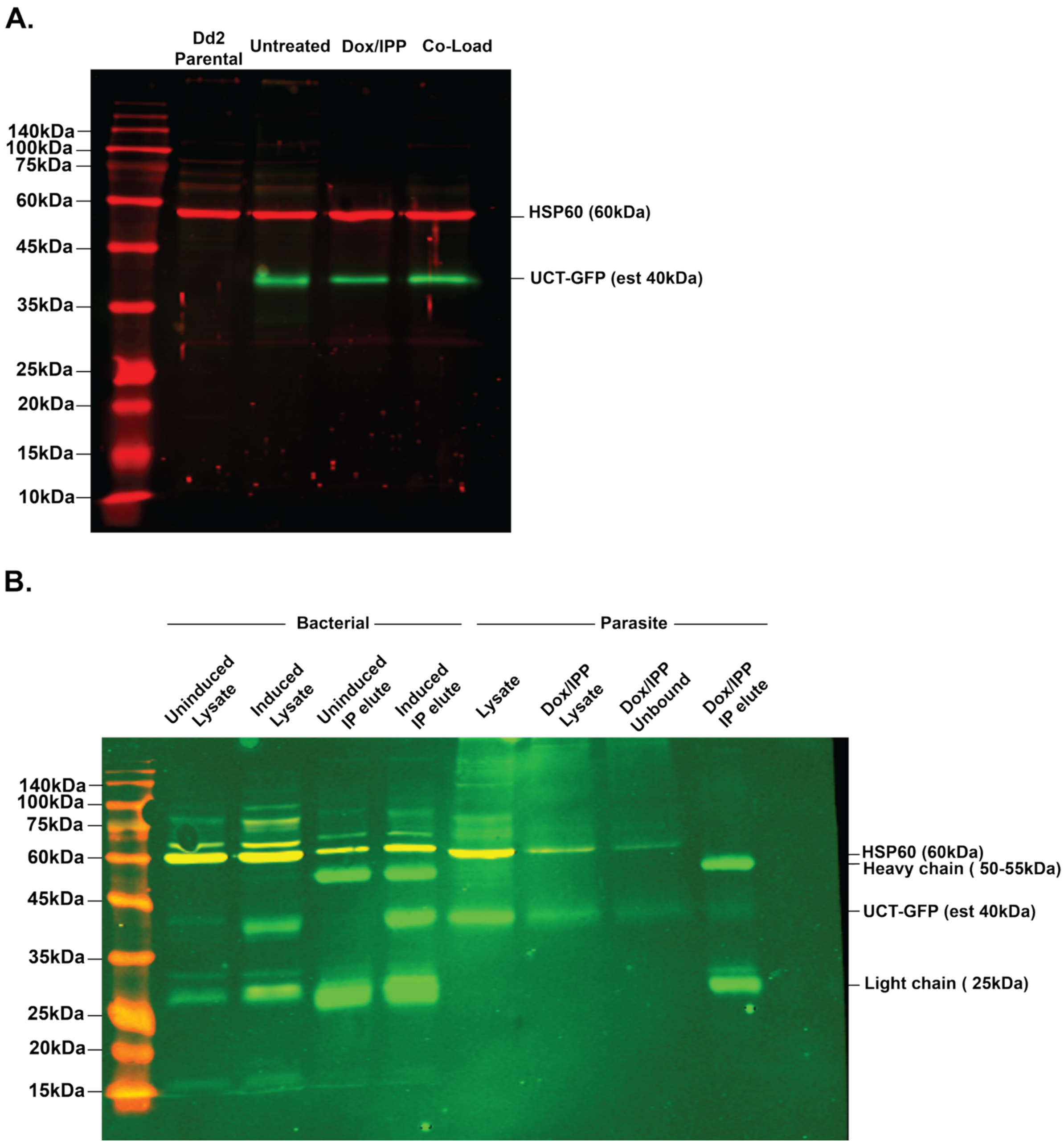


**Figure S4. Labeled western blots showing** (A) parental Dd2, untreated UCT-GFP, dox/IPP treated UCT-GFP, and co-loaded untreated and dox/IPP parasites stained with anti-GFP and anti-HSP60 antibodies (B) anti-GFP immunoprecipitation of UCT-GFP in E. *coli* and parasites stained with anti-GFP and anti-HSP60 antibodies.

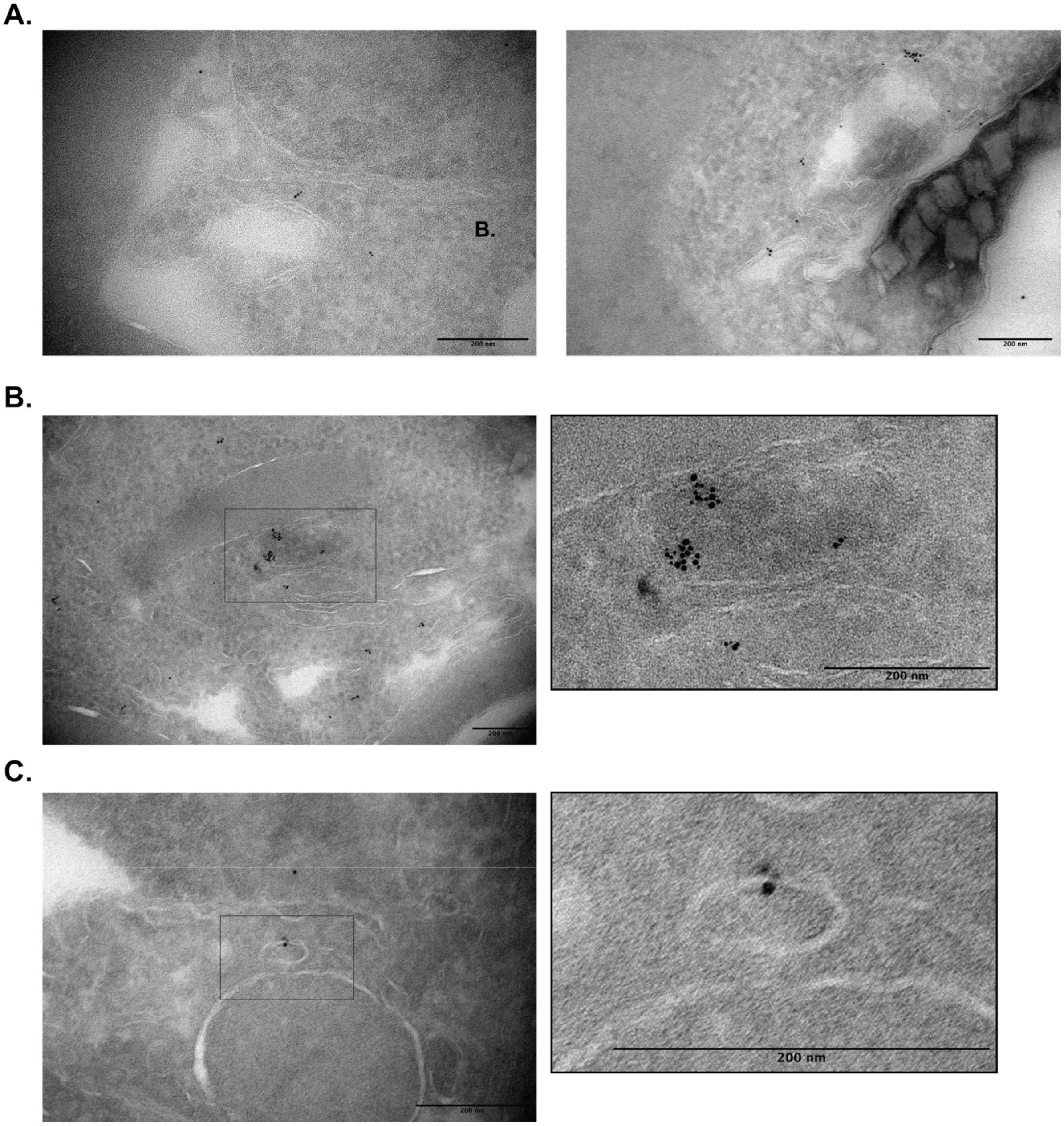


**Figure S5. UCT localizes to the membranous regions of the apicoplast.** Transmission electron microscopy images of (A) uncropped main figure images and (B) and (C) further examples of anti-GFP and 5nM colloidal gold antibodies marking UCT-GFP at the membranous regions of the apicoplast, black boxes indicate the zoomed image.

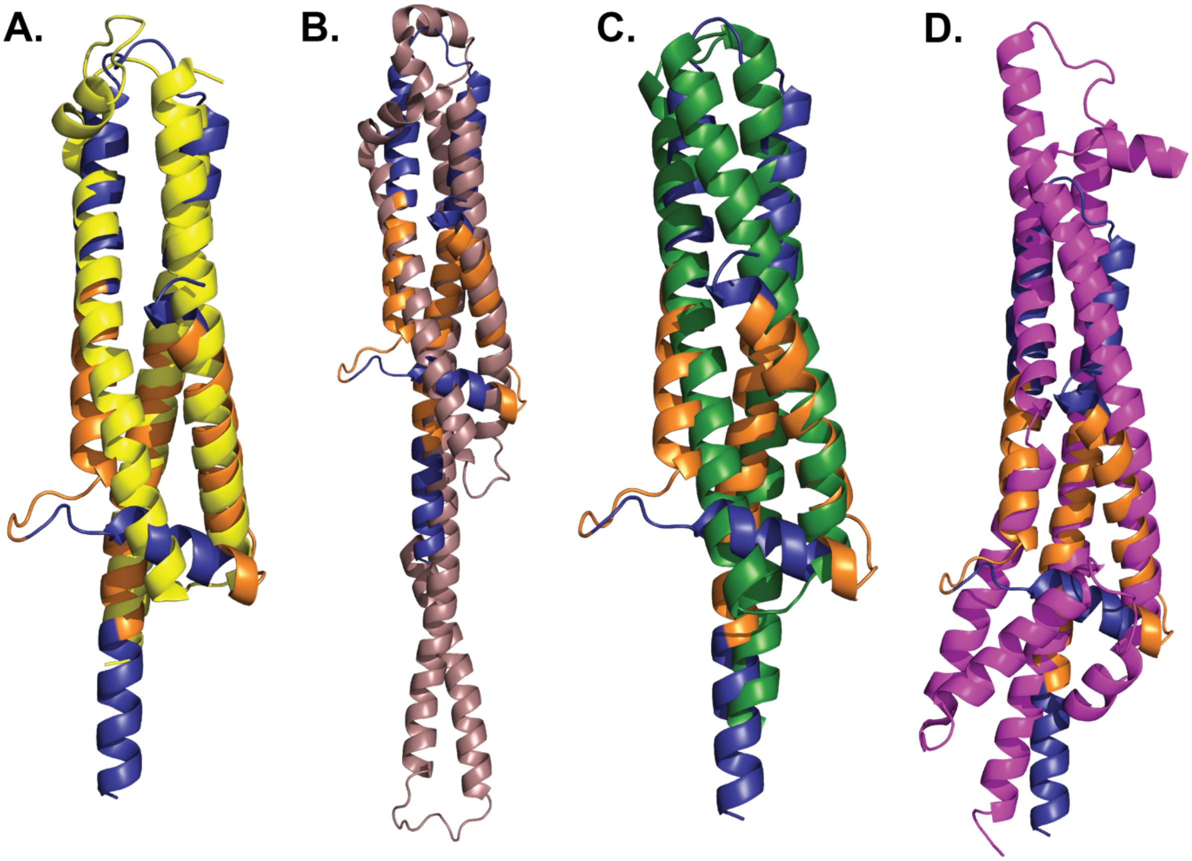


**E.**

| | CorA Yeast | CorA Human | CorA *E. coli* | ExbB |
|---|---|---|---|---|
| **Z-score** | 8.0 | 8.2 | 8.9 | 8.9 |
| **RMSD** | 2.8 | 4.1 | 3.0 | 3.5 |

**Figure S6.** UCT has structural similarity to known metal transporters, including (A) the yeast CorA magnesium transporter (PDB 3RKG) lacking the N-terminal globular domain, (B) the human CorA magnesium transporter (PDB 8IP4 lacking the N-terminal globular domain, (C) the *E. coli* CorA magnesium transporter (PDB 5N9Y) lacking the N-terminal globular domain, and (D) the *Cyanobacterial* ExbB transporter (PDB 5SV1). (E) Table showing Z-score and RMSD (Å) values obtained from the DALI server search.

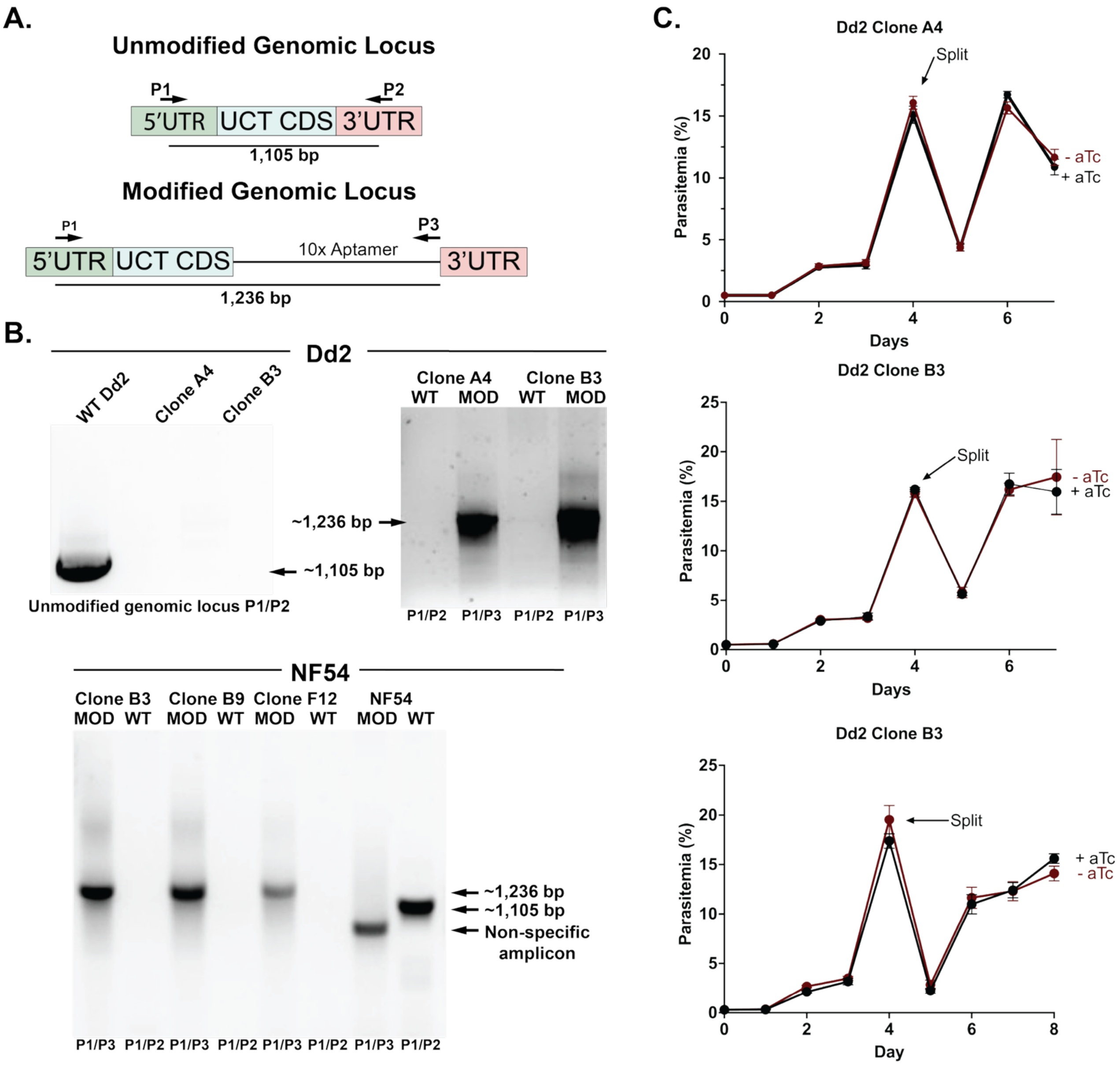


**Figure S7. Tagging UCT at the genomic locus for conditional knockdown** (A) Schematic representation of unmodified (WT) or modified (mod) UCT genomic locus, indicating primer sets used to identify each locus. (B) Agarose gels of PCR amplicons showing successful modification of the UCT genomic locus in Dd2 clones A4 and B3 and NF54 clones B3, B9, and F12 *P. falciparum* parasites. (C) Growth assays of UCT KD Dd2 clonal lines A4 and B3. Each growth assay was performed in biological triplicate, and cultures were split on day 4.

A.

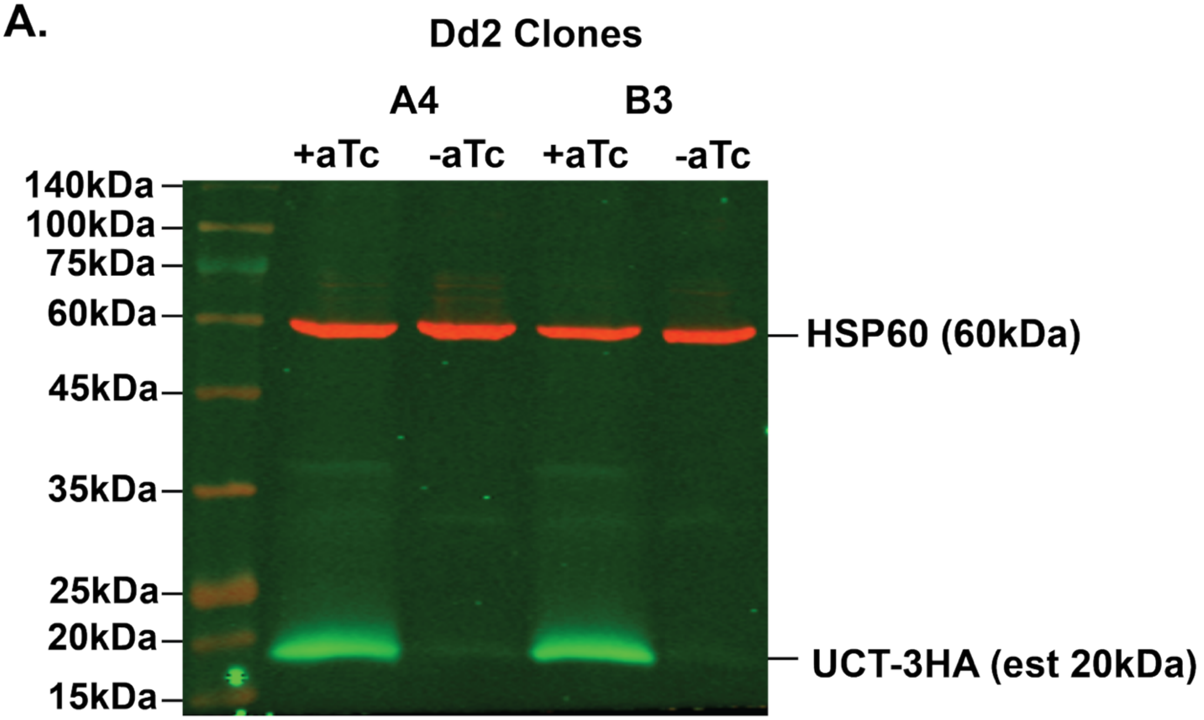


B.

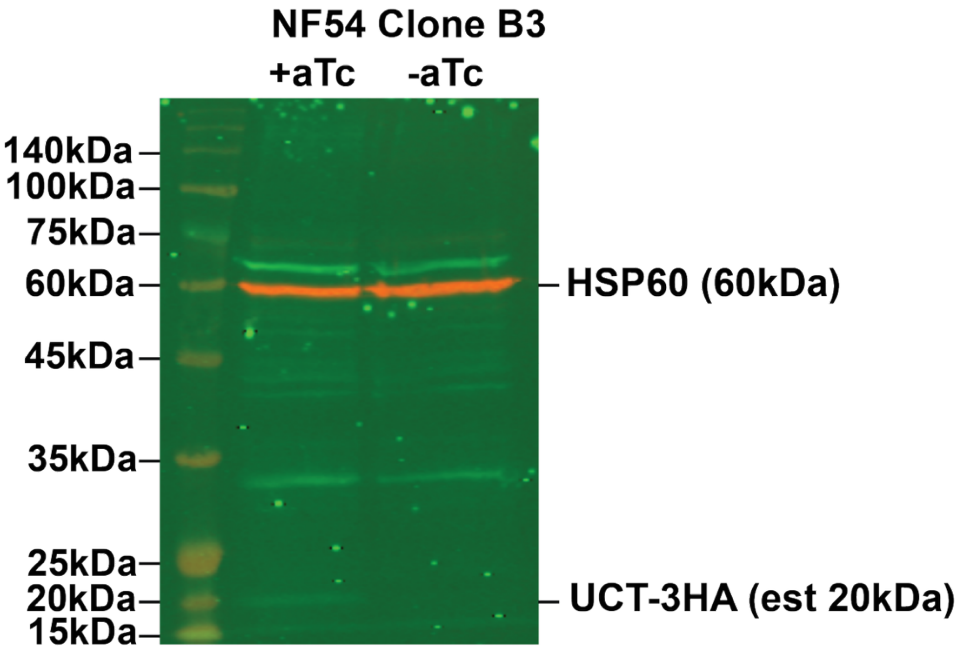


**Figure S8. Full SDS-PAGE western blot analysis of UCT knockdown.** Western blot analysis of endogenously tagged UCT-3HA in (A) Dd2 clones A4 and B3 and (B) NF54 clone B3 ± aTc stained with anti-HA and anti-HSP60 antibodies.

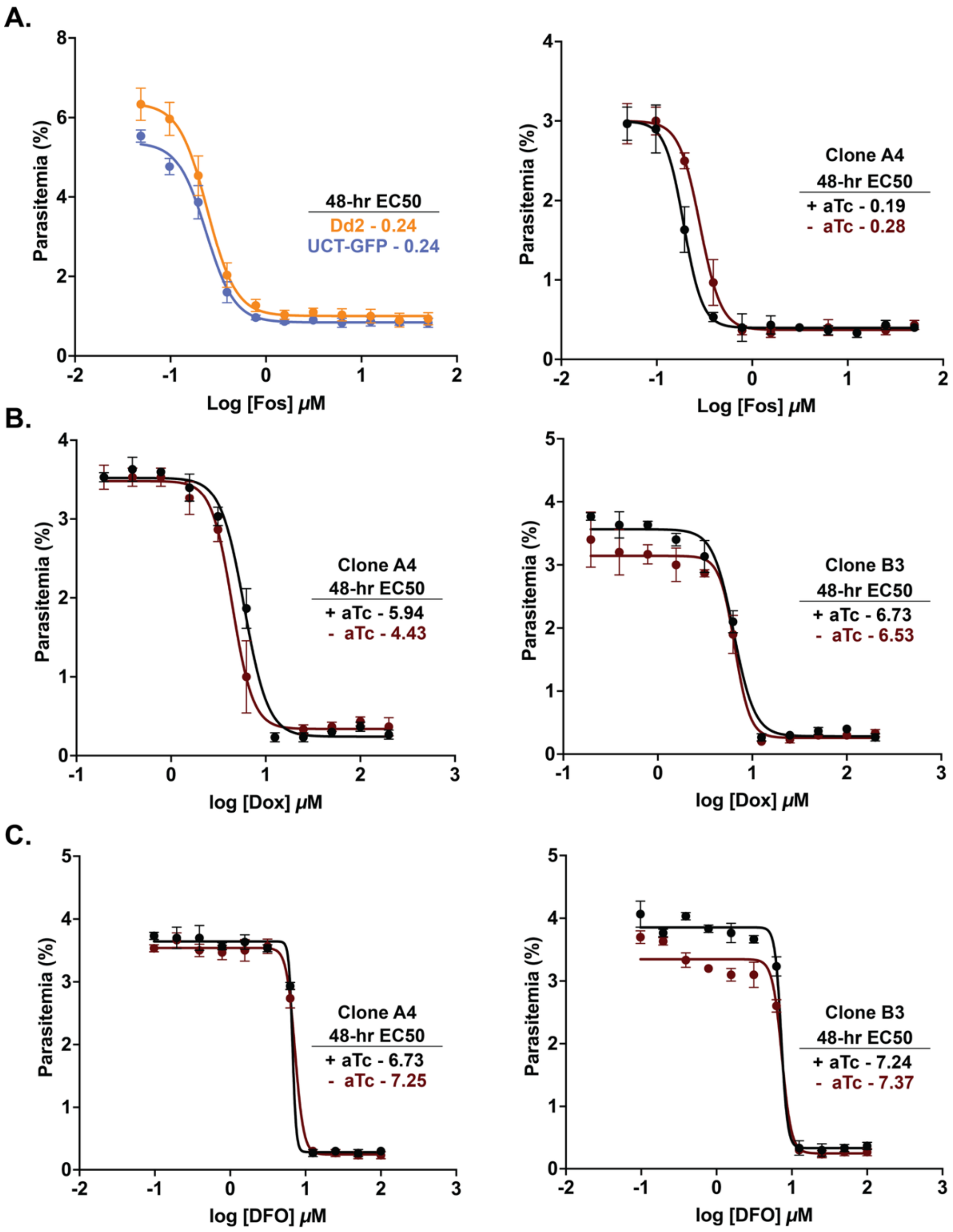


**Figure S9. Determination of 48-hour $EC_{50}$ values.** Dose-response curves of parasite growth versus (A) fosmidomycin (FOS) concentration using Dd2, UCT-GFP, and UCT KD parasites, (B) doxycycline (dox) concentration using UCT KD parasites, and (C) deferoxamine (DFO) concentration using UCT KD parasites. Data points and error bars show the average SD of biological triplicate measurements and fit with a four-parameter logistic dose-response model in GraphPad Prism.

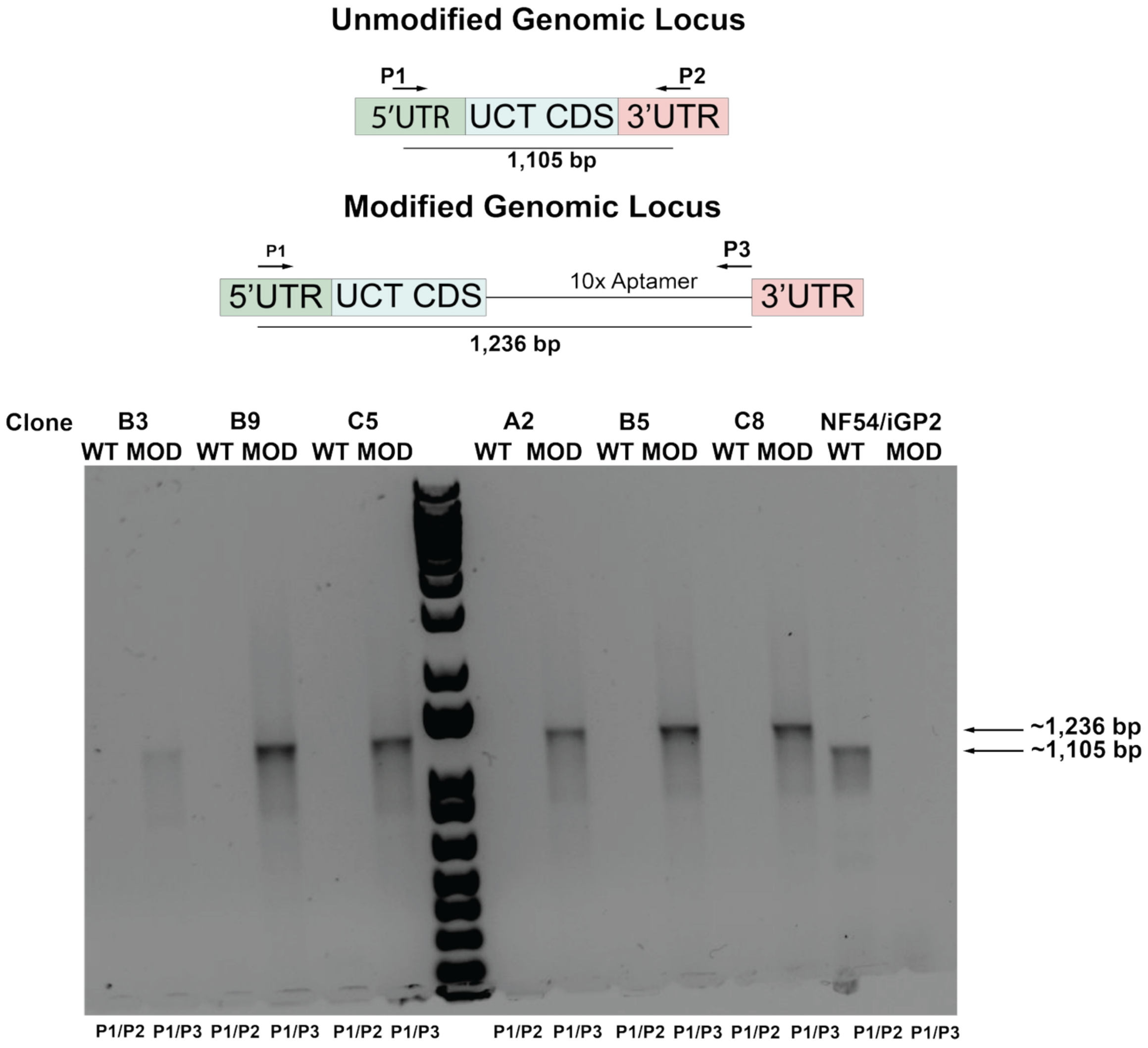


**Figure S10. Tagging UCT at the genomic locus for conditional knockdown in NF54/iGP2 parasites.** Schematic representation of unmodified (WT) or modified (mod) UCT genomic locus, indicating primer sets used to identify each locus and agarose gels of PCR amplicons showing successful modification of the UCT genomic locus in NF54/iGP2 clones B3, B9, C5, A2, B5, and C8 *P. falciparum* parasites.

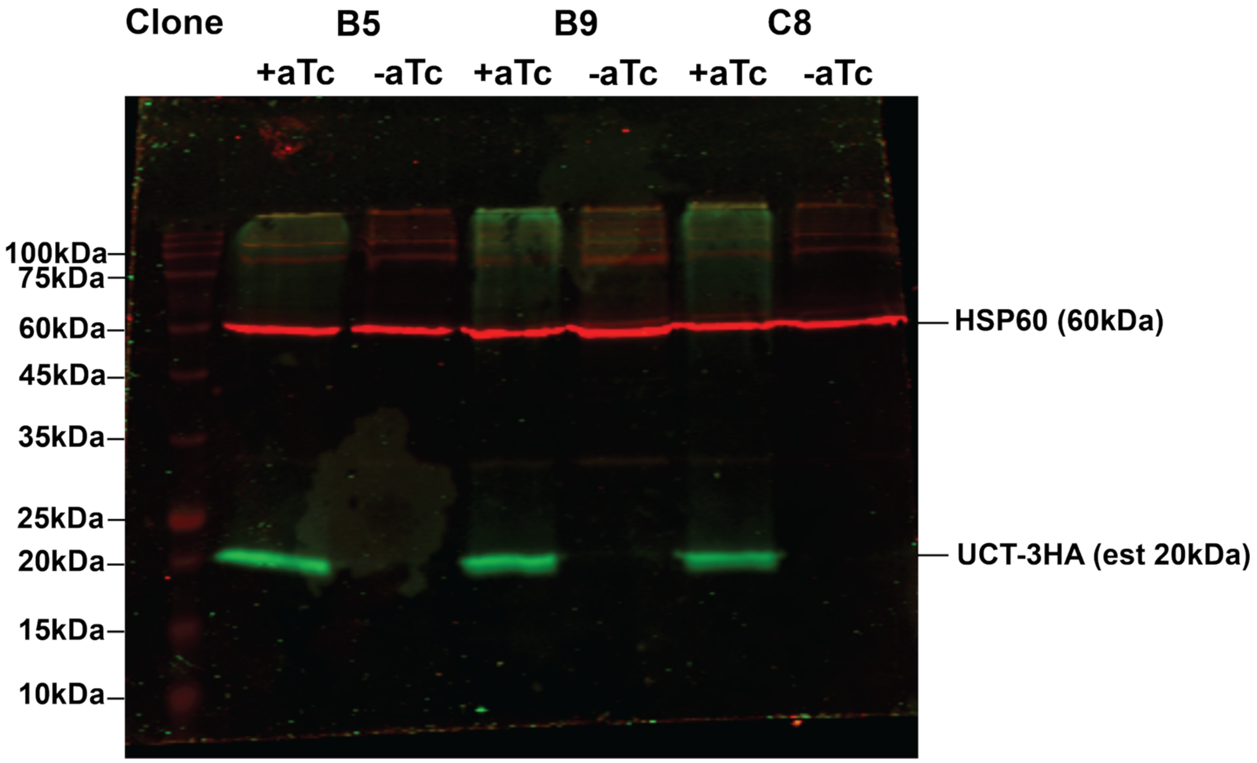


**Figure S11. Full SDS-PAGE western blot analysis of UCT knockdown.** Western blot analysis of endogenously tagged UCT-3HA in NF54/iGP2 clones B5, B9, and C8 in ± aTc, induced with glucosamine washout (-GlcN), and harvested on day 9 after induction without asexual arrest, then stained with anti-HA and anti-HSP60 antibodies.

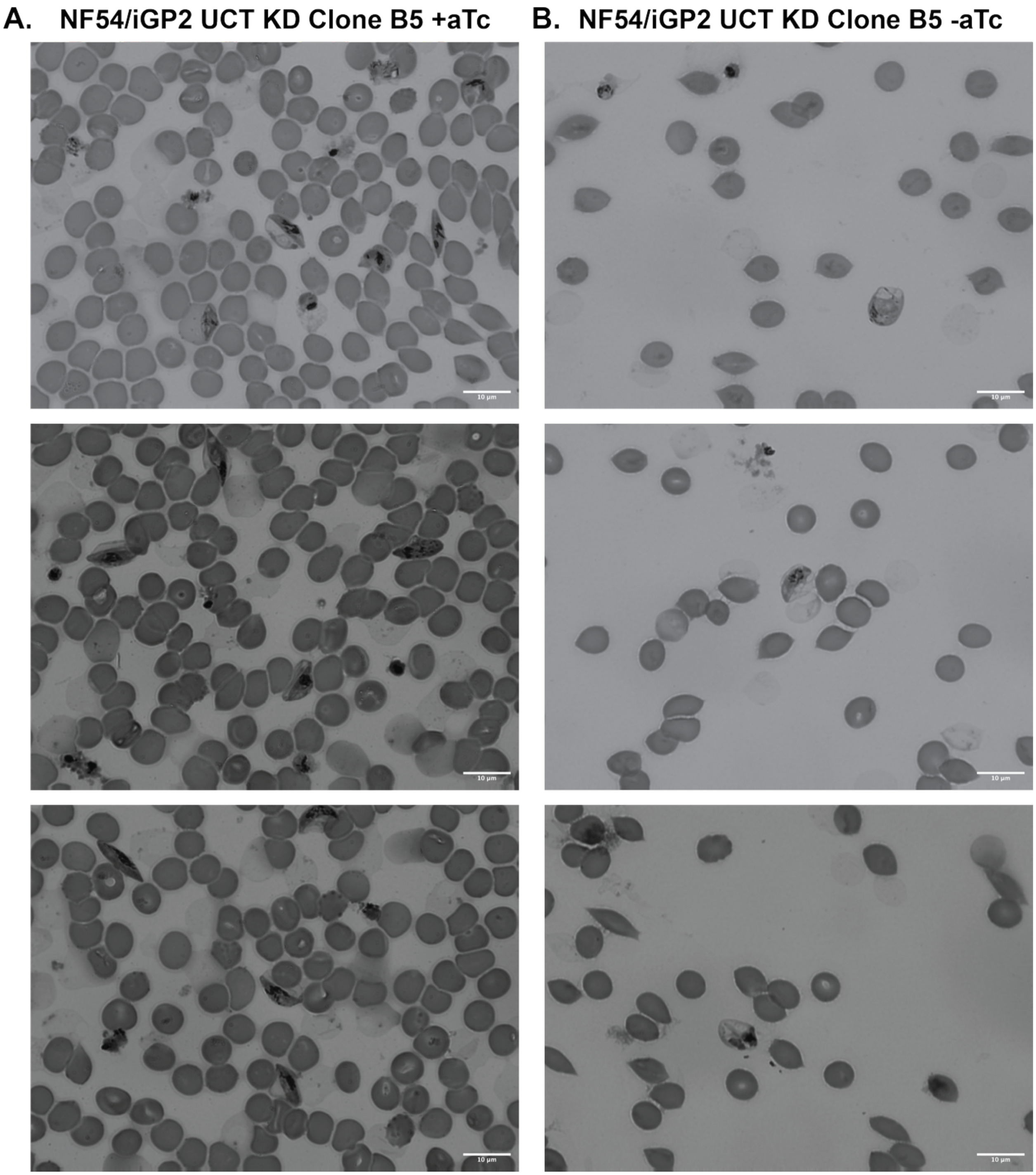


**Figure S12. Representative field images of NF54/iGP2 clone B5** in (A) +aTc and (B) -aTc conditions on day 9 after induction.

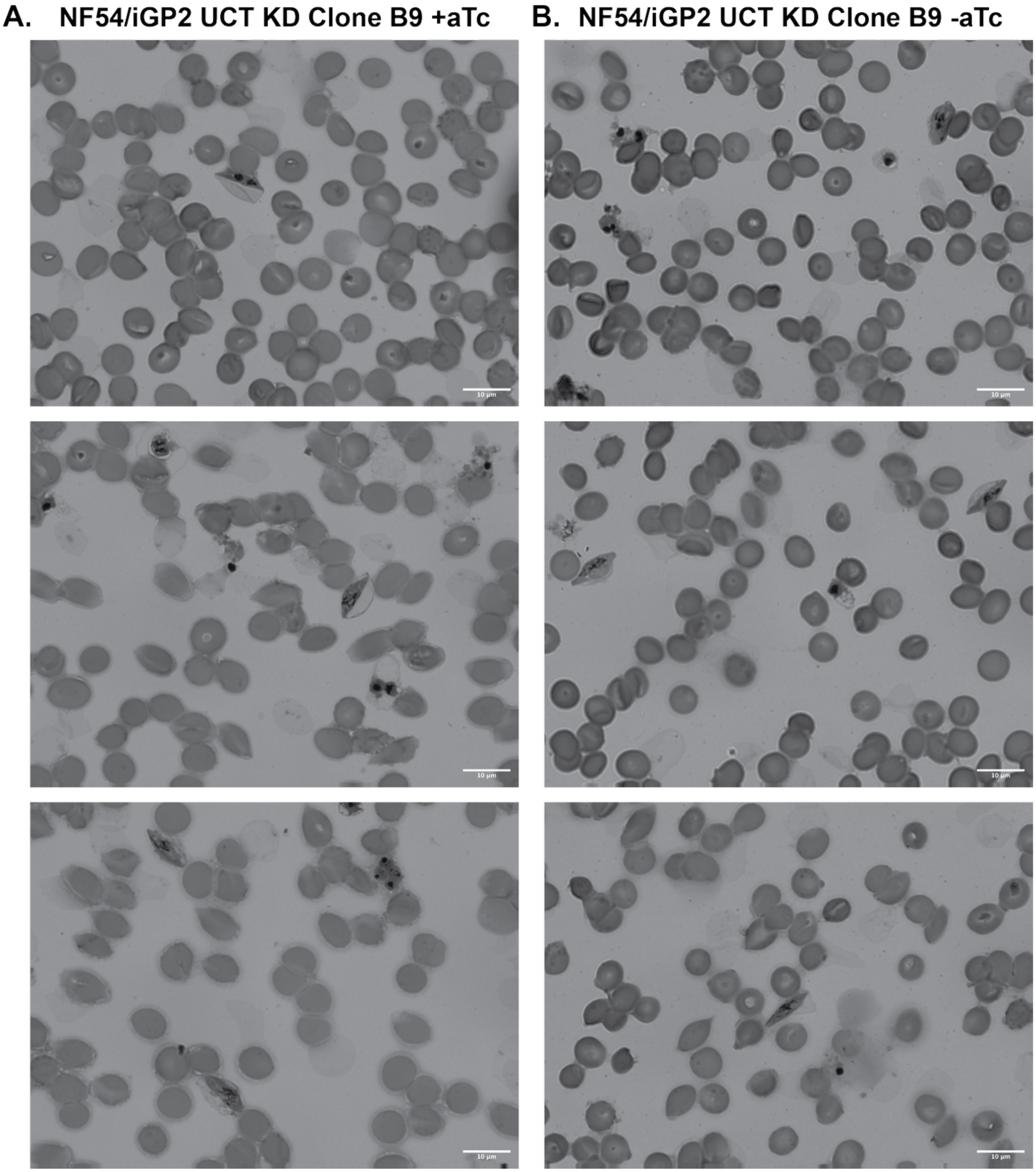


**Figure S13. Representative field images of NF54/iGP2 clone B9** in (A) +aTc and (B) -aTc conditions on day 9 after induction.

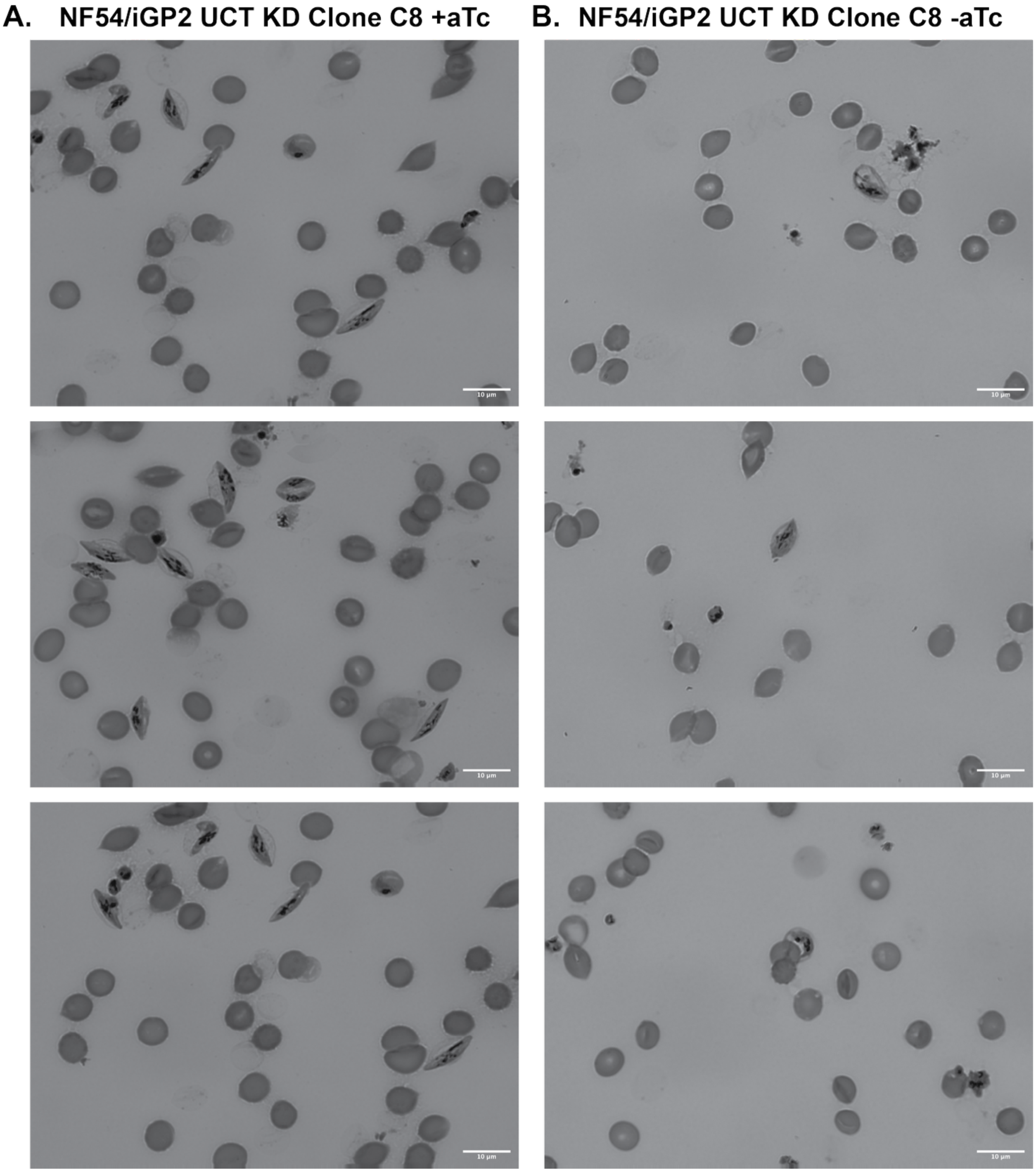


**Figure S14. Representative field images of NF54/iGP2 clone C8** in (A) +aTc and (B) -aTc conditions on day 9 after induction.

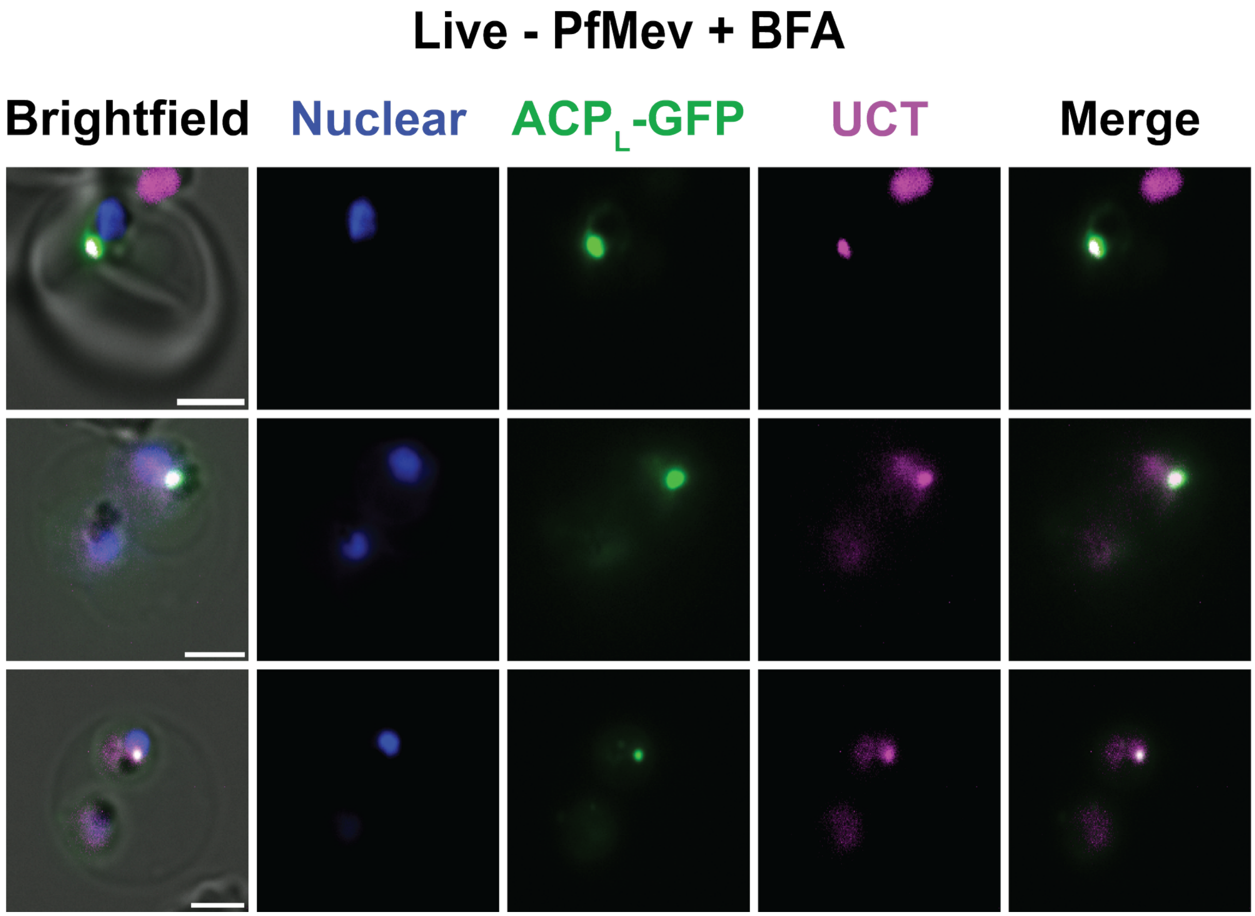


**Figure S15.** UCT localization is not sensitive to brefeldin A treatment. Live-parasite fluorescent microscopy of synchronized UCT-mS PfMev parasites treated with 5 µg/mL brefeldin A (BFA) and imaged after 18 hours. Scale bar = 2.5 µm.

| Primers used in this study | | |
|---|---|---|
| **No** | **Name** | **Sequence (5' to 3')** |
| 1 | UCT_TEOE_F | CGAATAAACACGATTTTTTCTCGAGATGACATTTTTAAAATTTATATATTTGATAGTTGTGCCCTTAGG |
| 2 | UCT_TEOE_GFP_F | TGCTGCATAATGTGCTGCACCTGGCCTAGGTAGGAGCTTTATTTGTCTTTTCCTTTGTTCT |
| 3 | UCT_NTERM_GFP_R | TGCTGCATAATGTGCTGCACCTGGCCTAGGACCAATCTTCTTTCTACAAAACGAATAACTAAACG |
| 4 | mScarlet_TEOE_F | AAATAAAGCTCCTACCTAGGGGAGGTGGCTCTGGCGGGGGATCAATGGTGAGCAAGGGC |
| 5 | mScarlet_TEOE_R | TATATAACTCGACGCCTAGGTTACTTGTACAGCTCGTCCA |
| 6 | UCT-Cas9CDSgRNA2-F | TCATATTAAGTAATAATATTAGTGCGAATGAGCTAATCACGTTTTAGAGCTAGAAATAGC |
| 7 | UCT-Cas9CDSgRNA2-R | GCTATTTCTAGCTCTAAAACGTGATTAGCTCATTCGCACTAATATTATTACTTAATATGA |
| 8 | UCT-Cas9CDSgRNA3-F | TCATATTAAGTAATAATATTTAGTGCGAATGAGCTAATCAGTTTTAGAGCTAGAAATAGC |
| 9 | UCT-Cas9CDSgRNA3-R | GCTATTTCTAGCTCTAAAACTGATTAGCTCATTCGCACTAAATATTATTACTTAATATGA |
| 10 | UCT-Cas9UTRgRNA5-F | TCATATTAAGTAATAATATTCAAATGATAAAATGTGTAGCGTTTTAGAGCTAGAAATAGC |
| 11 | UCT-Cas9UTRgRNA5-R | GCTATTTCTAGCTCTAAAACGCTACACATTTTATCATTTGAATATTATTACTTAATATGA |
| 12 | UCT_3UTR_KD_F | ATATATATCCAATGGCCCCTTTCCGGGCGCGCCTTTTTCATATACGTACACTTTTTAAATGTATGATGGATTATAGCT |
| 13 | UCT_3UTR_KD_R | AATATATAAATTTTAAAAATGTCATCTTAAGGTTTTCATATTTTATACTATGATTCACTTTGTTTTGTTTATATTTTCTTT |
| 14 | UCT_CDS_Frag1_F | GAATCATAGTATAAAATATGAAAACCTTAAGATGACATTTTTAAAATTTATATATTTGATAGTTGTGCCCTTAGG |
| 15 | UCT_CDS_Frag1_R | GTTATACCAGTAATGAGCTCATTTGCACTATAC |
| 16 | UCT_CDS_Frag2_F | AAATATGTATAGTGCAAATGAGCTCATTACTGGTATAAC |
| 17 | UCT_CDS_Frag2_R | CAGGTACGTCATAAGGGTATCCGGAGACGTCTAGGAGCTTTATTTGTCTTTTCCTTTGTTCTAAATATTTAACAT |
| 18 | Integration P1_F | CTGATAATTTATCGTATTTTCCTTTTG |
| 19 | Integration P2_R | GTTTTCATATTTTATACTATGATTCAC |
| 20 | Integration P3_R | CTAGACTAGGTTCCAAGATCTCCCG |
| 21 | UCT_pET24a_F | AAGAAGGAGATATACTCGAGATGACATTTTTAAAATTTATATATTTGATAGTTGTGCCCTTAGG |
| 22 | GFP_pET24a_R | TTGTTAGCAGCCGGATCTCACTTATTTGTACAGTTCATCCATGCCATGTGT |

Table S1. List of primer sets used.
